# Defining the ESKAPE pathogen prophage repertoire with PHORAGER

**DOI:** 10.64898/2026.08.05.742953

**Authors:** Xena Dyball, Alise J Ponsero, James A D Docherty, Andrea Telatin, Emmanuelle H Crost, Nathalie Juge, Ryan Cook, Evelien M Adriaenssens

**Affiliations:** Quadram Institute Bioscience, Norwich Research Park, Norwich, NR4 7UQ, UK; Centre for Microbial Interactions, Norwich Research Park, Norwich, NR4 7UG, UK; University of East Anglia, Norwich, NR4 7UG, UK

## Abstract

Prophages are major drivers of bacterial evolution, mediating horizontal gene transfer and lysogenic conversion to alter host phenotypes. Nevertheless, identifying prophages within bacterial genomes remains challenging due to their heterogeneity and similarity to other mobile genetic elements. Here we present PHORAGER (Prophage Hunting, vOtu Retrieval, Annotation and Genomic ExploRation), a scalable Nextflow pipeline for the standardised identification and quality assessment of prophages from bacterial genomes. PHORAGER incorporates bacterial genome pre-processing, consolidation of predictions from multiple mining tools, annotation-based filtering to reduce false positives, and generation of ready-to-analyse summary tables. We validated PHORAGER using 30,824 publicly available ESKAPE pathogen genomes. PHORAGER recovered more high-quality prophages than individual mining tools alone, and through extensive quality assessments removed a substantial number of false-positive predictions. In total 23,132 putative prophages were identified, the majority belonging to the class *Caudoviricetes*, and exhibiting a high degree of host-specificity. Putative antimicrobial resistance genes were detected in 0.48% of prophages, whereas virulence factors were most abundant in *S. aureus* prophages. ESKAPE prophages also frequently encoded anti-phage defence systems. PHORAGER is freely available as open-source software and the ESKAPE prophage collection generated in this study provides a reusable resource for further investigations.

**GRAPHICAL ABTRACT:** 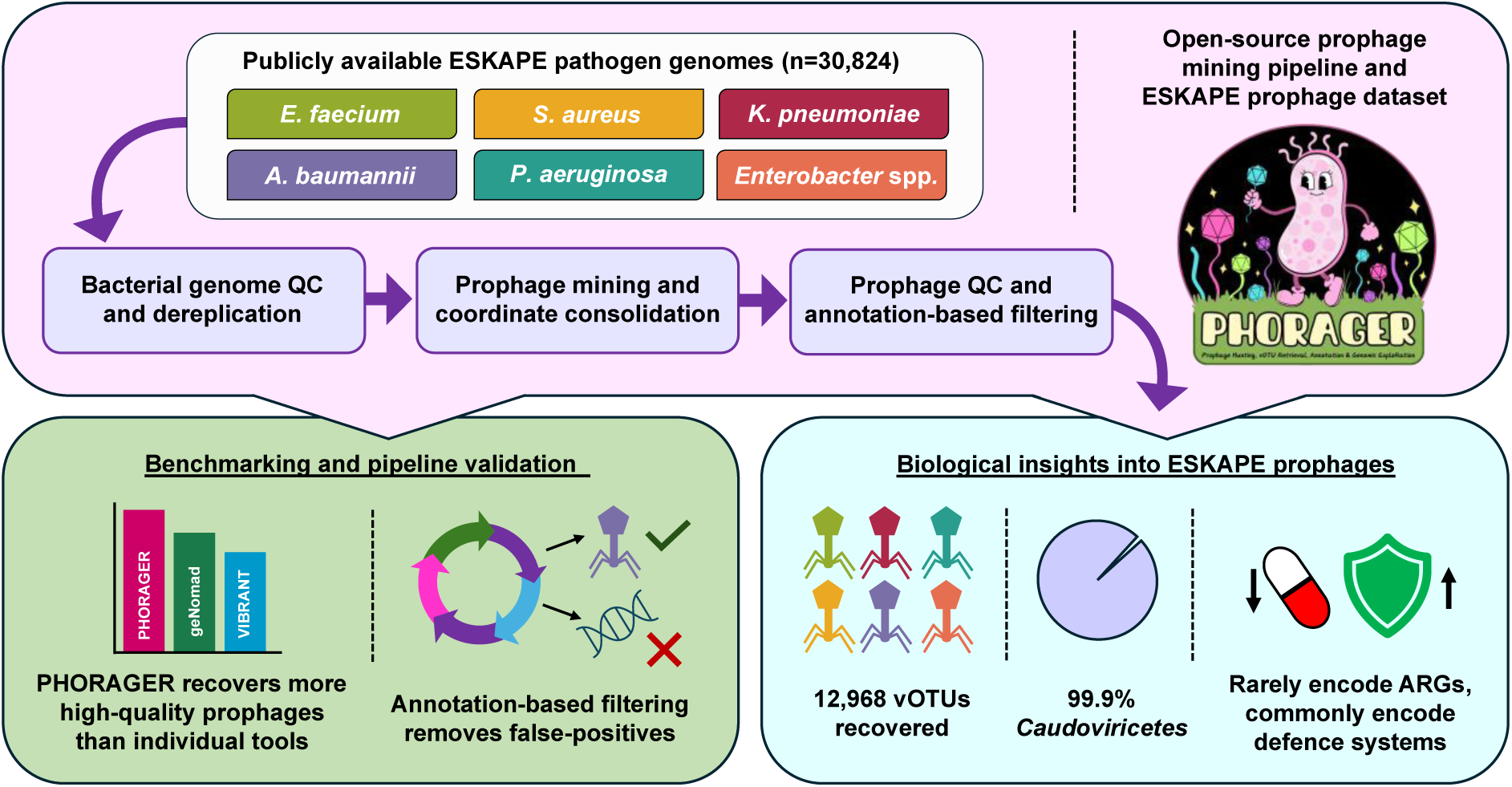

## INTRODUCTION

Bacteriophages (phages) are the natural viral predators of bacteria and are widely regarded as the most abundant biological entities on Earth (1,2). They play important roles in microbial ecology and evolution, as they can modulate bacterial populations and drive genomic diversity through infection (3). Phages mainly replicate via the lytic and lysogenic cycles, the latter of which involves integration of the viral genetic material into the bacterial chromosome to form a prophage. Prophages may persist indefinitely within their lysogenic host and spread vertically within populations via bacterial replication, alternatively they may switch to the lytic cycle and rapidly produce viral particles if prompted by an induction trigger (4,5). Prophages provide novel genes to their bacterial host, including toxins, auxiliary viral genes (AVGs), and defence systems, which can be expressed independently of the phage infection cycle and alter the microbial phenotype through lysogenic conversion (6–9). For instance, the ESKAPE pathogens – *Enterococcus faecium*, *Staphylococcus aureus*, *Klebsiella pneumoniae*, *Acinetobacter baumannii*, *Pseudomonas aeruginosa* and *Enterobacter* spp., a group of clinically important bacterial species and genera identified as emerging threats to public health, can attribute some of their clinical persistence to prophage-encoded genes such as toxins, immune evasion mechanisms, and enhanced biofilm formation (10–13). Prophages therefore have influence over a range of cellular processes and are responsible for much of the strain-level variation we see among bacterial populations (14).

Despite their significance, prophages can be challenging to isolate and study in the laboratory setting, which is why bioinformatics-based approaches are popular among researchers (15). The rapid development of high-throughput next-generation sequencing technologies and subsequent availability of bacterial genomes in public databases has additionally helped accelerate bioinformatic-based mining efforts (16). As such the estimate of the number of bacteria that harbour prophages (lysogens) has continued to increase overtime, with one study suggesting that 93% of bacteria encode at least one prophage (17). Nonetheless, bioinformatic prediction of prophages still has its limitations, largely due to the heterogeneity of phage genomes and lack of universal marker gene to link them (18). Their similarity to the surrounding bacterial genes, resemblance to other types of mobile genetic elements, such as transposons, and potential degradation over time, can additionally hinder prophage identification (19,20).

To overcome some of these issues, different prophage mining tools have been developed which employ a diverse set of methods to optimise prophage identification from bacterial sequencing data. To date there are over 90 virus mining tools available, with over 50 specialising in prophage detection (https://github.com/shandley/awesome-virome). While many of these tools were developed within the last decade (e.g. PHASTER and PHASTEST), a few of them have been in use for almost 20 years, including Phage_Finder and Prophinder, (21–24). The earlier tools relied largely on sequence similarity-based methods to identify prophages, comparing nucleotide or protein sequences to custom databases of known phages or phage genes (24). Although sequence similarity searching works well, it may prevent the discovery of more novel prophages that are unlike those present in the current databases. Therefore, many of the newer tools have transitioned to using updated mining approaches or a combination of methods, such as machine learning models and biological metrics, including GC content, gene length/direction, and k-mer signatures (25–27). The extensive range of mining tools currently available offers a remarkable selection, however navigating this diversity can pose challenges when determining which one to use.

At present there is no standardised workflow for prophage mining, so researchers use a mixture of tools and typically implement limited quality checks on their putative prophages before performing downstream analysis. Due to these differing workflows, the accuracy of results generated can vary greatly between studies. Here we present PHORAGER (Prophage Hunting, vOtu Retrieval, Annotation and Genomic ExploRation) (https://github.com/aponsero/PHORAGER), a scalable Nextflow prophage mining pipeline for the standardised identification of prophages from bacterial genomes. The pipeline includes robust bacterial genome pre-processing steps, methods to de-replicate predictions generated by multiple miners, and advanced quality assessments to reduce the number of false positives identified.

To validate PHORAGER, we tested it on a dataset of 30,824 publicly available ESKAPE pathogen genomes. The ESKAPE pathogens were selected because they represent a panel of clinically important bacteria with broad research interest globally. These pathogens also include Gram-positive and Gram-negative organisms spanning a range of genome sizes, all of which have been sequenced extensively, allowing us to evaluate PHORAGER’s ability to recover prophages from a diverse collection of bacterial genomes.

## MATERIALS AND METHODS

### Pipeline Design

PHORAGER comprises four main workflows (Figure 1), which can be run consecutively or separately depending on the user’s requirements.

1. Bacterial workflow: Quality assessment and dereplication of bacterial genomes
2. Prophage workflow: Prediction and consolidation of putative prophage sequences
3. Annotation workflow: Quality assessment, annotation-based filtering, and dereplication of prophage sequences
4. Summary workflow: Generation of bacterial genome and prophage summary tables

**Figure 1:**
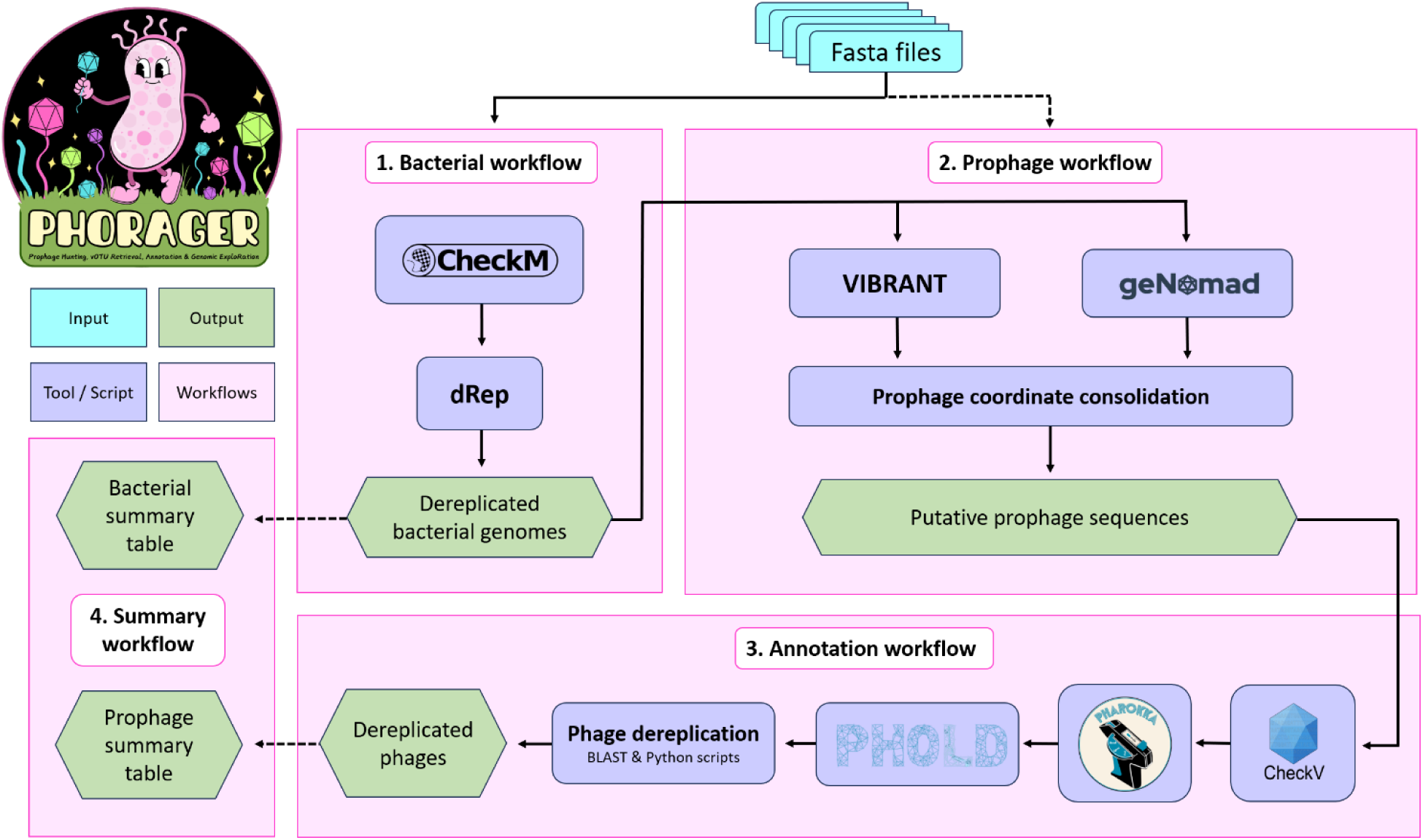
PHORAGER workflow. Overall design of the pipeline, depicting the four main workflows, bacterial pre-processing, prophage mining, prophage processing, and summary table generation, in addition to the key tools/steps involved at each stage.

### Bacterial Workflow

To begin, PHORAGER employs CheckM2 v1.0.1 to assess the quality of the bacterial genomes provided by the user (28–31). The pipeline applies a user-customisable quality cut-off to filter the genomes based on their completeness and contamination scores (default parameters = ≥95% completeness and <5% contamination). Genomes that satisfy the quality requirements are carried forward in the pipeline, and those that fail are discarded.

Genomes are then dereplicated using dRep v3.5.0 (32). Genome dereplication is an important step in the pipeline as it helps to remove redundancy from the user’s dataset while still maintaining the genomic diversity. The threshold at which the genomes are dereplicated can be modified based on the user’s preference, however the default parameters dereplicate the sequences at the approximate strain level (99.9% ANI) (33).

### Prophage Workflow

Following bacterial preprocessing, the pipeline enters the prophage mining stage. Using the dereplicated genomes, the pipeline operates two prophage mining tools, VIBRANT (Virus Identification By iteRative ANnoTation) v1.2.1 and geNomad v1.8.1, in parallel to identify putative prophage sequences (26,27). These tools were selected as they are unique in their approaches to prophage prediction, thus providing greater opportunity for a more diverse set of prophages to be identified.

As PHORAGER employs two miners it is likely that some prophages will be predicted more than once. In these cases, it is important to remove repeated predictions, nevertheless it can be difficult to decide which prediction is correct, especially when each tool has identified different boundaries for the prophage. To overcome this issue PHORAGER uses a custom python script to obtain the genomic coordinates of each predicted prophage, detect overlapping coordinate ranges, and determine the greatest range by identifying the earliest start and latest end coordinate (Supplementary figure 1). PHORAGER then uses the consolidated coordinates to extract the prophage sequences from the bacterial genomes and compile them in preparation for processing.

### Annotation Workflow

Initial processing of the prophages begins with CheckV v1.0.3, which assesses the quality of the viral genomes by detecting and removing bacterial contamination, measuring genome completeness, and identifying closed genomes (34). PHORAGER runs CheckV’s ‘end-to-end’ workflow to obtain the quality score for each prophage. The user is then able to filter the results and remove those that do not meet a specified quality or length requirement, for example using the default parameters prophages that are categorised as low-quality or ‘not determined’ and are <5 kb in length are discarded.

The filtered prophage sequences are then annotated using pharokka v1.9.1, which is a specialised bacteriophage annotation tool (35). Pharokka is run with the default settings, using PHANOTATE for gene prediction and the standard PHROG, CARD, and VFDB databases to assign functional annotations (36–44). Additionally, the tool Phold v0.2.0, a novel phage annotation tool based on protein structural homology, can be run following Pharokka to improve the number of functional annotations per sequence (40,45–48). The annotations are then parsed to determine the percentage of structural genes, e.g. connector, head and packaging, or tail genes, present in each sequence. The prophage sequences can then be filtered based on their structural gene percentages, with default parameters removing sequences with a Pharokka percentage <5 and a Phold percentage <10.

The remaining sequences can then be dereplicated to ensure the output only contains unique prophages. The viral dereplication process involves using blast v2.16.0 and the custom python scripts provided by CheckV, which work in combination to group the prophages into clusters based on pairwise ANI (34). Users can define the clustering parameters. However, it is recommended to dereplicate at the approximate species level – 95% ANI and 85% aligned fraction (AF) (15).

### Summary workflow

PHORAGER also has a summarise function, allowing the user to generate summary tables in tab-delimited (.tsv) format for easy downstream analysis. A bacterial genome summary table can be generated which comprises quality control (QC) metrics and prophage counts per genome, additionally a prophage summary table can be generated which contains information about each prophage’s bacterial host, cluster, QC score, length, and CDS count.

### Pipeline implementation

PHORAGER is implemented as a Nextflow DSL2 workflow fronted by a Python command-line wrapper (49). The wrapper exposes the pipeline through argparse subcommands (config, install, bacterial, prophage, annotation, summarise); each subcommand validates user input and then constructs and dispatches a single Nextflow run main.nf --workflow <name> invocation, so that all execution, environment provisioning, and data staging are delegated to Nextflow. Full documentation is available on GitHub (https://github.com/aponsero/PHORAGER/wiki) and WorkflowHub (https://workflowhub.eu/workflows/2211).

### Reusability

The pipeline follows a modular structure in which each bioinformatics tool and each custom processing step is encapsulated as a self-contained Nextflow process (a module), and the workflows compose these modules through DSL2 include statements. This separation allows individual steps to be reused across workflows and recombined without code duplication. Pipeline configuration is likewise factored into discrete, included files: global defaults and resource directives, per-process software specifications, execution profiles, and tool/database metadata are each maintained separately and merged at runtime. Tool versions, container image identifiers, and database descriptors are declared centrally as structured parameters (container_specs, database_specs), giving a single authoritative source for these definitions. The summary toolkit extends this principle further through a registry pattern, so that a new summary table type can be added by creating a single module and registering it, without modifying the dispatch logic.

### Scalability

PHORAGER exploits Nextflow’s dataflow model to parallelise across both samples and sequences. Input genomes enter the pipeline as independent items on a channel, and the prophage-mining processes fan out to process each genome concurrently, with VIBRANT and geNomad executing in parallel as independent branches of the same input channel. In the annotation workflow, the consolidated prophage set is split into individual sequences that are then annotated independently, allowing per-prophage parallelism across the annotation and structural-prediction steps. Computational resources are allocated per process through named directives, with memory requests configured to scale on automatic retry and a bounded retry strategy applied to transient failures; concurrency for the most resource-intensive tools is capped to avoid oversubscription. Because resource requests and scheduling are expressed through Nextflow’s executor abstraction rather than hard-coded, the same workflow definition can run unchanged from a single workstation up to a multi-node compute environment.

### Portability

The pipeline supports two interchangeable software backends: (1) Conda/Bioconda and (2) Singularity, selected at runtime through execution profiles, with Singularity as the default (50,51). Each process defines both execution paths and branches on the active profile, so that no change to the workflow code is required to switch environments. Software versions are pinned identically across the Conda specifications and the container image tags to keep the two backends consistent and the analyses reproducible; containers are sourced from public registries (BioContainers and, for custom environments, Sylabs Cloud) and are pulled and cached automatically on first use (52).

### Computational time and resources benchmark

To characterise the computational cost of PHORAGER, we benchmarked the prophage and annotation workflows on a set of 100 bacterial genomes taken randomly from the HumGut database (53). The bacterial workflow was excluded, as its runtime is governed chiefly by dRep and its dereplication settings rather than by the prophage-mining logic. Prophage detection was benchmarked in three configurations 1) geNomad only, 2) VIBRANT only, and 3) both tools combined, and annotation in two: 1) full detailed annotation (Pharokka followed by Phold, CPU-only) versus 2) detailed annotation disabled; the two annotation configurations were run against an identical prophage set so that only the annotation path varied. Each configuration was run in triplicate, sequentially using the Singularity backend, with a preceding warm-up run to eliminate cold-cache effects. All benchmark runs were executed on a single compute node equipped with one 84-core AMD EPYC 9634 processor (single socket, one thread per core, 84 logical CPUs) and 510,000 MB of RAM (≈498 GiB), running Linux (kernel 5.14) under SLURM 23.02.8. Each benchmark job was allocated 16 CPUs and 200 GB of memory, and the full sweep completed in 3 d 2 h 34 min of wall-clock time. Per-process resource use was captured from Nextflow’s built-in execution trace (raw output), from which wall-clock time, CPU-hours (execution time × allocated CPUs), and peak resident memory (peak RSS) were derived. Only successfully completed tasks were included in the aggregation. The computational resources used by each step is detailed in supplementary tables 1-3, and where dependent on the included tools in each sub-workflow.

### Dataset Curation

All *Enterococcus faecium, Staphylococcus aureus, Acinetobacter baumannii, Pseudomonas aeruginosa*, and *Enterobacter* spp. genomes that were complete at the contig level were downloaded from the National Centre for Biotechnology Information (NCBI) RefSeq database on 10^th^ October 2025 in fasta format. Due to the substantially higher number of available *Klebsiella pneumoniae* genomes, those that were complete at the chromosome level were downloaded from the NCBI database on 22nd October 2025.

### Running PHORAGER

PHORAGER (version v0.5.0) was run on the Norwich Bioscience Institute high-performance computing cluster using the default parameters.

### Analysis of PHORAGER summary files

The.tsv summary tables and.txt summaries generated by PHORAGER were parsed and analysed in R using dplyr, tidyr, and tidyverse (54–56). All data visualisation was performed using the R packages ggplot2, ggrastr, ggside, eulerr, and ggpubr, except for the Sankey plot which was produced using SankeyMATIC (SankeyMATIC: Make Beautiful Flow Diagrams) (57–61).

### Taxonomic analysis of prophages with vConTACT3

vConTACT3 (v3.1.3) was run using the --proteins option (62). For this, fasta amino acid (.faa) files were generated from the Phold GenBank files for the final dereplicated prophage dataset. Additionally, a genes2genome and genome lengths file in parquet format were also generated and used as input for vConTACT3. To allow for the network to be opened in Cytoscape v3.10.4, the network was filtered for edges with distance ≤ 0.5 and shared genes ≥ 2 (63). This reduced the number of edges from ∼76 million to ∼2 million prior to visualisation. This filtering applies only to the visualisation and does not affect the taxonomic assignments made by vConTACT3. Singletons are not displayed in the network.

### Detection of putative AMR genes and virulence factors using ABRicate

ABRicate (https://github.com/tseemann/abricate) (v1.4.0) was run using the ResFinder and VFDB databases to generate output files in tab format (43,64,65). A custom python script was used to parse the ABRicate output and generate a network tsv file and node attributes tsv file that were visualised in Cytoscape (v3.10.4)(63).

### Identification and analysis of anti-phage defence systems

Coding sequences on the dereplicated bacterial genomes were predicted using Prodigal v2.6.3 and used as input for DefenseFinder v3.0.0 with model version 3.1.0, which utilises MacSyFinder v2 in its implementation (30,66,67). DefenseFinder was run with default parameters. Each system reported by DefenseFinder was treated as a single detected defence system. For each genome, the defence repertoire was summarised as the total number of systems and as the set of systems present, classified by DefenseFinder subtype.

Defence systems were classified as prophage-encoded or non-prophage by comparing the coordinates of each system’s constituent genes with the coordinates of the final prophage sequences from the same genome. A system was classified as prophage-encoded if at least half of its genes fell within a prophage region, and as non-prophage otherwise. Non-prophage systems were used for all analyses of host defence carriage, and prophage-encoded systems were summarised separately, including by the vConTACT3 family of the prophage in which they were located.

Statistical analyses were performed in R (v4.2.2). For each pathogen, Pearson’s correlation coefficient was used to assess the relationship between bacterial genome length and non-prophage defence-system count, and between genome length and prophage count. To test whether non-prophage defence-system count was associated with prophage count independently of genome length, a first-order partial correlation between the two was calculated for each pathogen, controlling for genome length. To identify individual defence subtypes associated with prophage carriage, each subtype present in 5 to 95% of a pathogen’s genomes was tested using a Poisson generalised linear model of prophage count, with subtype presence as the predictor and genome length (log-transformed) and the number of other non-prophage defence systems in the genome as covariates; heteroscedasticity-consistent (HC1) standard errors were used. Within each pathogen, p-values were corrected for multiple testing using the Benjamini-Hochberg procedure, and subtypes with an adjusted p-value below 0.05 were considered significant.

## RESULTS

### Creating a test dataset of ESKAPE pathogen genomes

To evaluate the performance of PHORAGER, we used it to detect prophages in a dataset of publicly available ESKAPE pathogen genomes. Collectively, the ESKAPE pathogens comprise a mix of Gram-positive and Gram-negative organisms that share several clinically significant characteristics, including the ability to thrive in healthcare environments, develop diverse antimicrobial resistance (AMR) mechanisms, and spread successfully on a global scale, making them an interesting case study for prophage analysis (13).

A total of 30,824 ESKAPE pathogen genomes were used as input for PHORAGER. The dataset comprised 2,594 *E. faecium*, 11,102 *S. aureus*, 3,582 *K. pneumoniae*, 4,776 *A. baumannii*, 5,272 *P. aeruginosa*, and 3,498 *Enterobacter* spp. genomes obtained from NCBI (Figure 2A). The median genome lengths were 2.96 Mb, 2.83 Mb, 5.67 Mb, 3.96 Mb, 6.74 Mb, and 4.95 Mb for *E. faecium, S. aureus*, *K. pneumoniae*, *A. baumannii*, *P. aeruginosa*, and *Enterobacter* spp., respectively (Supplementary Figure 2).

**Figure 2:**
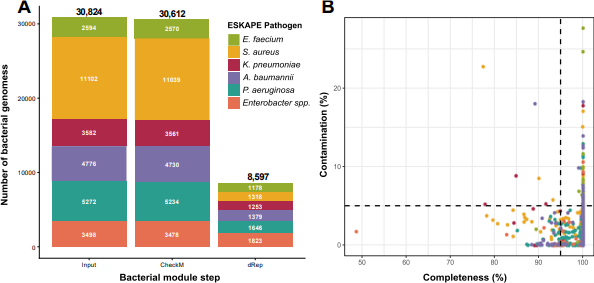
Bacterial genome QC. **(A)** The number of ESKAPE pathogen genomes that passed each filtering step in the PHORAGER bacterial workflow. (**B)** The CheckM2 completeness and contamination scores for each bacterial genome. The black dotted lines indicate the default filtering thresholds used by PHORAGER of >95% completeness and <5% contamination. The lower right quadrant shows the genomes that passed CheckM2 filtering and were used as input for dRep.

### Sequence dereplication reveals a high degree of redundancy among publicly available ESKAPE pathogen genomes

Following execution of the bacterial module of PHORAGER, fewer than 1% of genomes for each pathogen failed initial quality filtering criteria (>95% completeness and <5% contamination) with CheckM2 and were removed from the dataset (Figure 2B). The remaining genomes were subsequently dereplicated at the strain level (99.9% ANI) using dRep.

Dereplication substantially reduced the number of genomes retained for each pathogen. *S. aureus* exhibited the greatest reduction, with 88% of the CheckM2-filtered genomes removed, indicating a high degree of redundancy among publicly available genomes in NCBI. In contrast, *Enterobacter* spp. showed the smallest reduction following dereplication (47%), suggesting that this group was the most genomically diverse among the ESKAPE pathogens, likely reflecting its classification as a genus rather than a single species.

The final dereplicated dataset comprised 1,178 *E. faecium*, 1,318 *S. aureus*, 1,253 *K. pneumoniae*, 1,379 *A. baumannii*, 1,646 *P. aeruginosa*, and 1,823 *Enterobacter* spp. genomes (Figure 2A, Supplementary Table 4).

### There is substantial overlap in the putative prophage sequences predicted by VIBRANT and geNomad

The dereplicated genomes were used as input for PHORAGER’s prophage mining module, which incorporates the tools geNomad and VIBRANT to identify viral sequences within the bacterial genomes. Across the ESKAPE dataset, geNomad predicted 53,542 putative prophages, while VIBRANT predicted 43,536 (Figure 3A). *Enterobacter* spp. generated the highest number of predictions from both tools, with 12,482 putative prophages identified by geNomad and 10,969 by VIBRANT.

**Figure 3:**
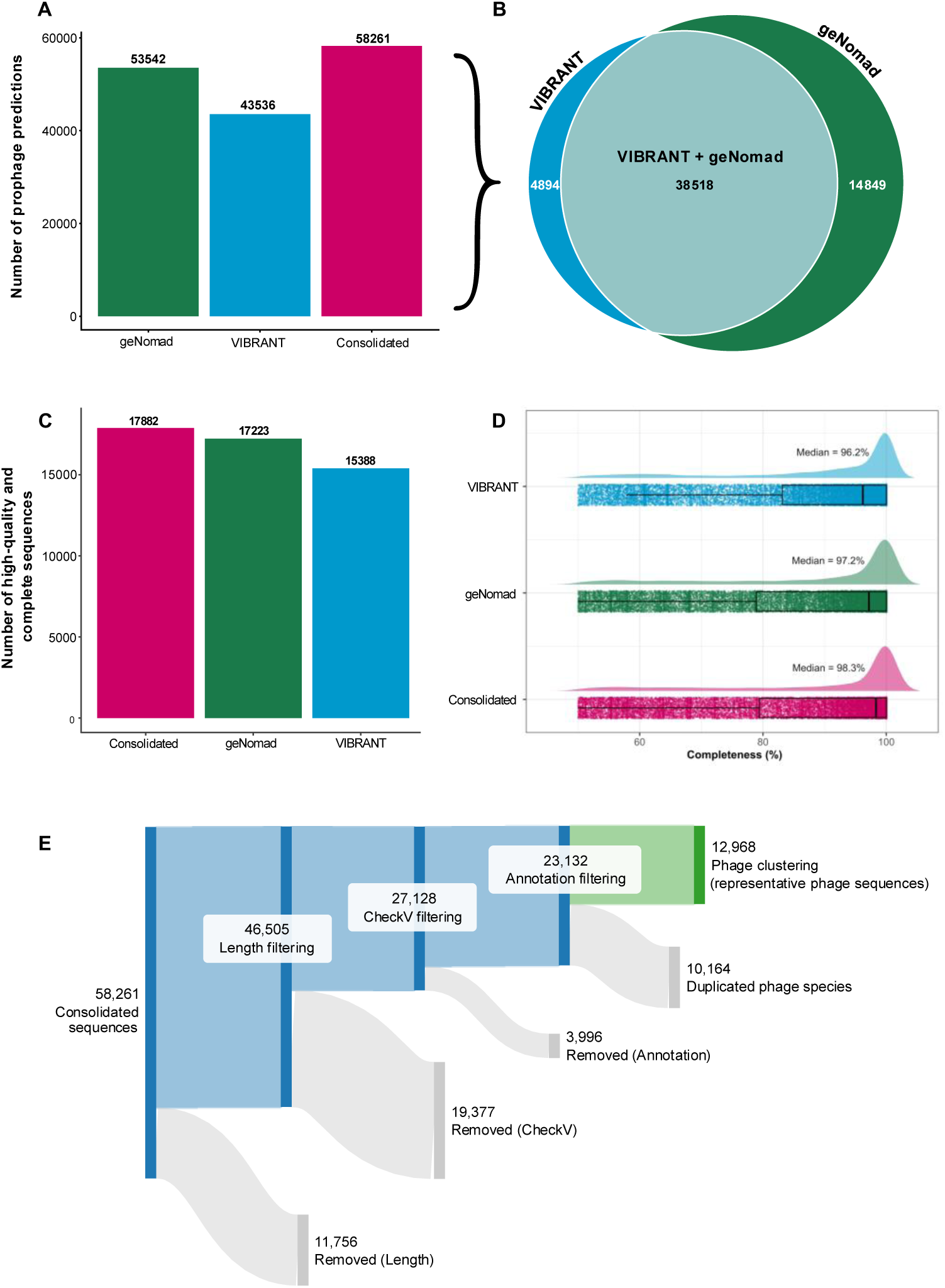
Coordinate consolidation removes substantial sequence redundancy and improves prophage quality and completeness. **(A)** Number of putative prophage sequences predicted by geNomad and VIBRANT, and the number of unique consolidated sequences produced when overlapping predictions from the two tools are merged. **(B)** The tool of origin for all the putative prophage sequences present in the consolidated dataset. **(C)** The number putative prophage sequences assigned ‘high-quality’ or ‘complete’ by CheckV for the individual prophage mining tool predictions versus the consolidated sequences. **(D)** The checkV completeness scores for the putative prophage sequences assigned ‘medium-quality’, ‘high-quality’, and ‘complete’. **(E)** Number of prophage sequences that passed each filtering stage in PHORAGER’s annotation workflow, starting with the consolidated sequences generated by the prophage workflow.

Overlapping prophage predictions generated by geNomad and VIBRANT were merged to produce consolidated sequences, reducing redundancy within the dataset. Approximately 66% of predictions generated by the two tools overlapped and were therefore consolidated. Of the individual tools, geNomad produced the highest proportion of unique prophage predictions (25%), compared with VIBRANT (8%) (Figure 3B, Supplementary Table 5).

Analysis of the consolidation patterns revealed that most consolidated sequences were generated by merging only two predictions, although one case involved the merging of nine individual predictions of smaller fragments. In all instances where two sequences were merged, the consolidated sequence comprised one prediction from geNomad and one from VIBRANT. For consolidated sequences generated from three predictions, 58% consisted of two geNomad predictions and one VIBRANT prediction, and the remaining 42% showed the reverse combination. In the single case where nine predictions were merged, eight originated from geNomad and one from VIBRANT (Supplementary Figure 3, Supplementary Table 5). Manual inspection suggested that this merged sequence is indeed a single prophage, although it is unclear why its prediction was so fragmented.

Following this consolidation step, a total of 58,261 non-redundant putative prophage sequences remained (Figure 3A).

### Consolidation of putative prophage predictions from multiple tools recovers higher quality prophage sequences

To validate the rationale for consolidating the prophage predictions produced by geNomad and VIBRANT, we compared the CheckV quality scores obtained for predictions generated from the individual tools with those from the consolidated sequences. Overall, sequence consolidation produced the greatest number of high-quality and complete sequences (17,882), compared with geNomad (17,223) and VIBRANT (15,388) (Figure 3C).

When analysed individually, all ESKAPE pathogens except *S. aureus* showed the same pattern, with consolidation producing the highest number of high-quality and complete prophage sequences, followed by geNomad and then VIBRANT (Supplementary Figure 4). In contrast, geNomad produced slightly more high-quality and complete sequences for *S. aureus* (2,224) than the consolidated dataset (2,192).

Additionally, we found that sequence consolidation also improves the completeness of the putative prophages, producing a median completeness score of 98.3%, compared to 96.2% and 97.2% for VIBRANT and geNomad respectively (Figure 3D).

### Advanced quality control steps filtered out a substantial amount of mis-identified prophage sequences

The 58,261 consolidated prophage sequences were input into PHORAGER’s annotation module. Sequences <5 kb were removed, resulting in the exclusion of 11,756 (20.2%) sequences. The remaining sequences were then filtered based on CheckV quality scores, retaining those classified as ‘medium-quality’, ‘high-quality’, or ‘complete’, producing a dataset of 27,128 sequences (Figure 3E).

Following annotation with Pharokka and Phold, sequences were further filtered based on the number and proportion of structural genes encoded. This annotation-based filtering step removed an additional 3,996 sequences. Of the removed sequences, 66% were classified as ‘medium-quality’ by CheckV, while a small proportion (1.4%) were predicted as ‘complete’ (Supplementary Figure 5A). Further investigation revealed that these ‘complete’ predictions were based on the prediction of the presence of high-confidence direct terminal repeats (DTRs) (Supplementary Figure 5B). In addition, 34.5% of the removed ‘complete’ sequences carried the CheckV warning ‘no viral genes detected’, highlighting the importance of this annotation-based filtering step to reduce potential false positives.

The resulting filtered prophage sequences were then clustered at the vOTU level using a 95% ANI threshold across an 85% aligned fraction, yielding 12,968 unique phage populations across the ESKAPE pathogens (Figure 3E). *Enterobacter spp.* had the highest number of phage vOTUs (4,157), whereas *E. faecium* had the fewest (923) (Supplementary Figure 6). Cluster analysis showed that *P. aeruginosa* harboured the largest individual cluster, which consisted of 324 members, suggesting that this phage population is highly conserved among *P. aeruginosa* strains.

When linking the final prophage dataset back to their bacterial hosts we find that 93.6% of ESKAPE pathogens are lysogens, harbouring at least one prophage (Supplementary Table 4). The percentage of lysogens does vary slightly between the individual pathogens, with *E. faecium* having the lowest percentage (89.7%) and *K. pneumoniae* having the highest (97.2%). The median prophage genome length ranged from 36.5 kb in *E. faecium* to 44.6 kb in *S. aureus* across the ESKAPE panel, with the three largest (*S. aureus*, *A. baumannii* and *K. pneumoniae*) all close to 44 kb (Supplementary Figure 7, Supplementary Table 6). The ESKAPE prophage dataset is available on Zenodo (DOI:10.5281/zenodo.18328113).

### Prophage carriage rates correlate with bacterial genome length

A commonly cited hypothesis is that the number of prophages within bacterial genomes positively correlates with genome length. We tested this relationship in the ESKAPE pathogen dataset and found a statistically significant, moderate, positive correlation between bacterial genome length and prophage count in all six species (R ranging from 0.46 to 0.61, all p<0.001; Supplementary Figure 8).

A similar trend was observed when examining prophage burden across genomes. Organisms with smaller genomes generally exhibited lower prophage counts. For example, *E. faecium,* which has a median genome length of 2.96 Mb, harboured a maximum of 5 prophages per genome, whereas *P. aeruginosa,* with a median genome length of 6.74 Mb, had up to 17 prophages in a single genome (Supplementary Figure 9).

### ESKAPE prophages are predominantly dsDNA tailed phages

To evaluate the ability of PHORAGER to detect taxonomically diverse prophages, we analysed the taxonomy of the final prophage dataset using vConTACT3. The detected prophages were assigned to 5 viral classes: *Caudoviricetes*, *Malgrandaviricetes,* and three putatively novel classes, encompassing both double-stranded and single-stranded DNA viruses (Supplementary Table 7). Nevertheless, although PHORAGER detected prophages spanning five viral classes, the dataset was overwhelmingly dominated by *Caudoviricetes* (99.9%).

Further taxonomic analysis classified the prophages into 115 viral families and 2,250 viral genera. Of these, 14.8% of families and 6% of genera corresponded to known taxa, with the remainder representing putatively novel families and genera. The largest known viral family was *Peduoviridae*, a family of double-stranded DNA phages that contains the well characterised temperate *E. coli* phage P2. *Peduoviridae* also demonstrated the greatest amount of sharing between ESKAPE pathogens, with prophages from *A. baumannii* (192), *Enterobacter* spp. (1526), *K. pneumoniae* (795), and *P. aeruginosa* (519) assigned to this family.

Analysis of the predicted viral genera showed that 64% were host-specific, containing prophages from only a single ESKAPE pathogen (Figure 4). Among the genera shared between multiple ESKAPE pathogens, prophages from *Enterobacter* spp. and *K. pneumoniae* most frequently co-occurred, likely reflecting the close evolutionary relationship between these organisms, which both belong to the bacterial family *Enterobacteriaceae*.

**Figure 4:**
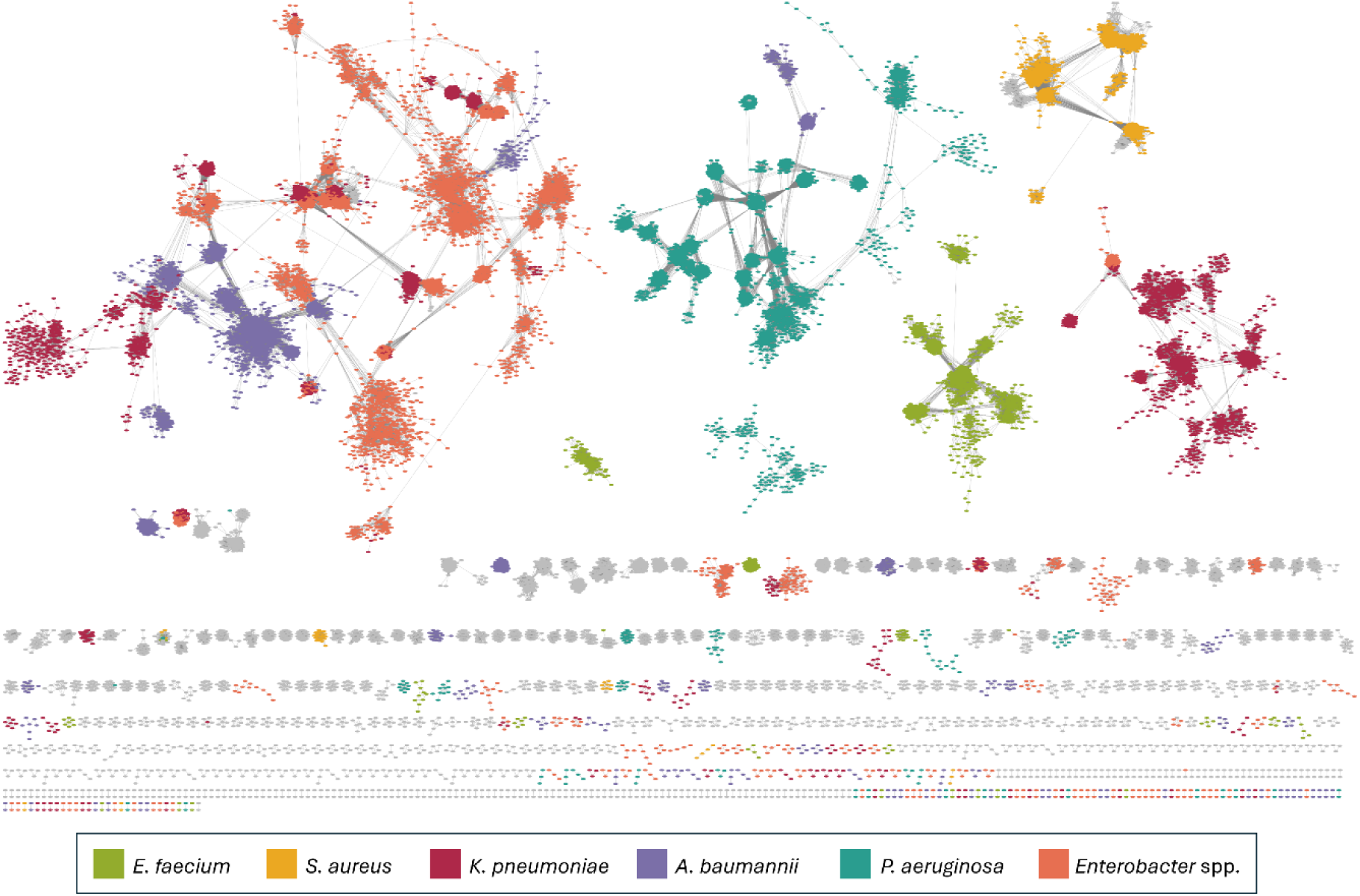
Taxonomic network of ESKAPE prophages. vConTACT 3 taxonomic network displaying the final prophage dataset. Each node represents a single prophage and is coloured based on their host organism. Grey nodes represent reference viruses from the vConTACT3 database. Singletons are not displayed.

### ESKAPE prophages encode low levels of putative AMR genes

To assess the potential contribution of the identified phages to bacterial pathogenicity, we used ABRicate to screen for AMR genes or virulence factors. Putative AMR genes (ARGs) were detected at very low frequencies, with only 0.48% of prophages across the ESKAPE panel encoding a predicted ARG. *A. baumannii* prophages encoded the greatest number of putative ARGs (n=24), whereas *P. aeruginosa* prophages encoded only a single putative ARG (Supplementary Table 8). Notably carbapenem resistance genes were exclusively predicted on *A. baumannii* prophages, and the majority of beta-lactam resistance genes were predicted in *K. pneumoniae* prophages (Figure 5A).

**Figure 5:**
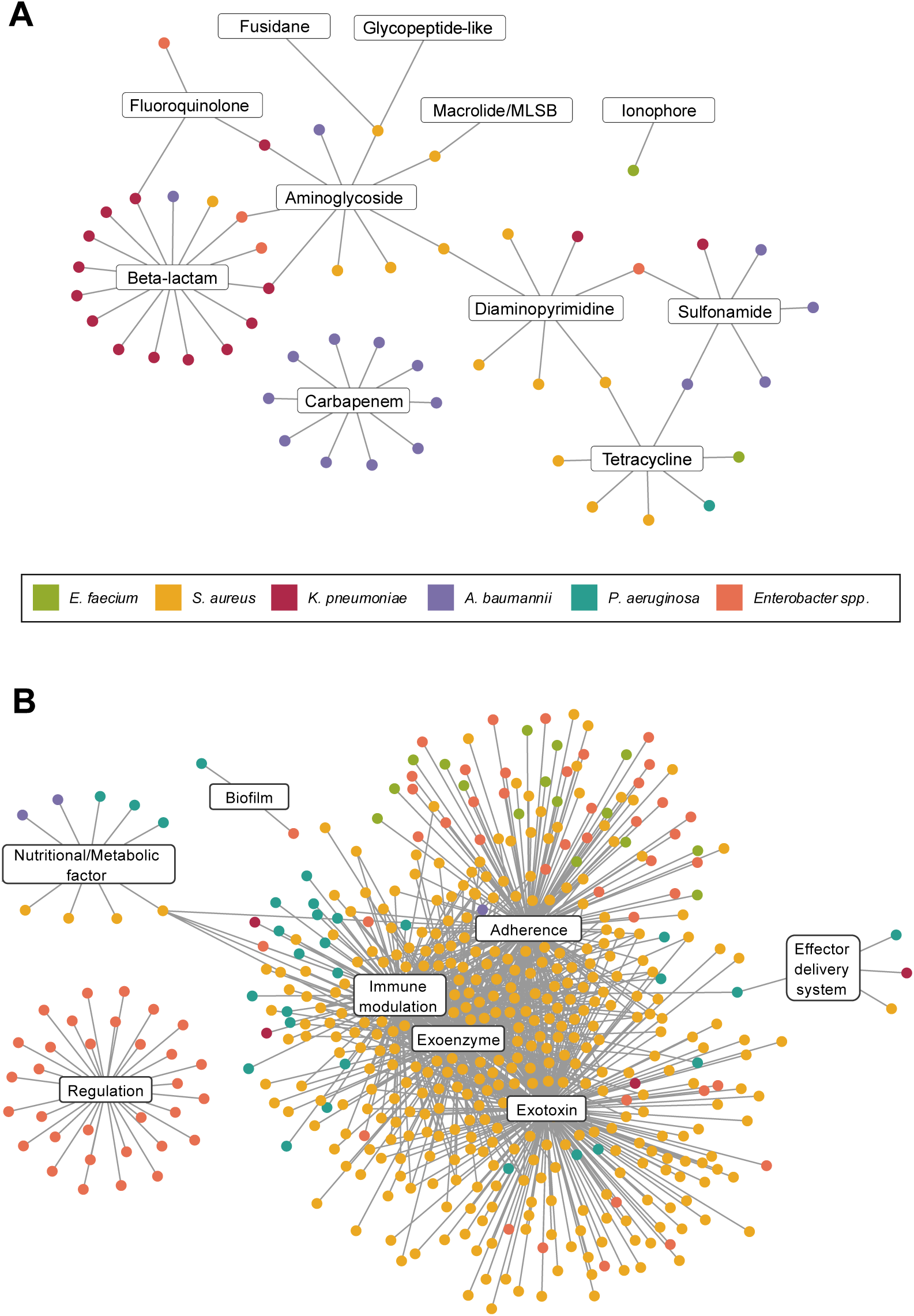
Putative prophage-encoded ARGs and virulence factors. **(A)** Network of ESKAPE prophages encoding antimicrobial resistance genes belonging to different antimicrobial classes, and **(B)** virulence factors grouped by VFDB category.

Virulence factors were also predicted across the ESKAPE pathogens, with *S. aureus* prophages encoding by far the largest number (1,021 genes). This was more than 100-fold higher than the second highest count, observed in *Enterobacter* spp. prophages (81 genes). Notably, 19% of the virulence factors encoded by *S. aureus* prophages corresponded to staphylokinase (*sak*), a well characterised *Staphylococcus*-specific exoenzyme associated with bacterial pathogenicity (Figure 5B). Interestingly, *A. baumannii* and *K. pneumoniae* prophages had the fewest virulence factors, with only three and four genes identified respectively, despite harbouring the highest numbers of putative ARGs (Supplementary Table 9).

### ESKAPE pathogens encode large and variable anti-phage defence repertoires

To examine the relationship between prophage carriage and anti-phage immunity, we identified defence systems in the 8,597 dereplicated ESKAPE genomes using DefenseFinder, detecting 86,321 systems in total (Supplementary Table 10). Almost all genomes encoded at least one system (98.8 to 100% across the six pathogens; Supplementary Figure 10). Repertoire size varied considerably between pathogens, being largest in *K. pneumoniae* (mean 14.9 systems per genome) and *P. aeruginosa* (13.6), smallest in *A. baumannii* (6.4) and *E. faecium* (6.0), and intermediate in *S. aureus* (8.5) and *Enterobacter* spp. (10.0). Repertoire diversity was independent of repertoire size. *Enterobacter* spp. encoded the greatest number of distinct subtypes (n=327), whereas *S. aureus* encoded the fewest (n=65). Type I restriction-modification was the most prevalent subtype in all six pathogens.

Defence systems were commonly encoded on prophages themselves, with 5,467 prophages (23.6%) encoding at least one (Figure 6A). The proportion of each pathogen’s total defence repertoire that was prophage-encoded ranged from 3.9% in *P. aeruginosa* to 12.4% in *A. baumannii*. Prophage-encoded defence was concentrated within particular viral lineages, with *Peduoviridae* the single largest contributor and several novel families (assigned by vConTACT3) encoding a defence system in more than 75% of their members. Because systems carried within prophages would otherwise confound any comparison of host defence with prophage carriage, we distinguished prophage-encoded systems from non-prophage systems and used only the non-prophage systems in the analyses that follow.

**Figure 6.**
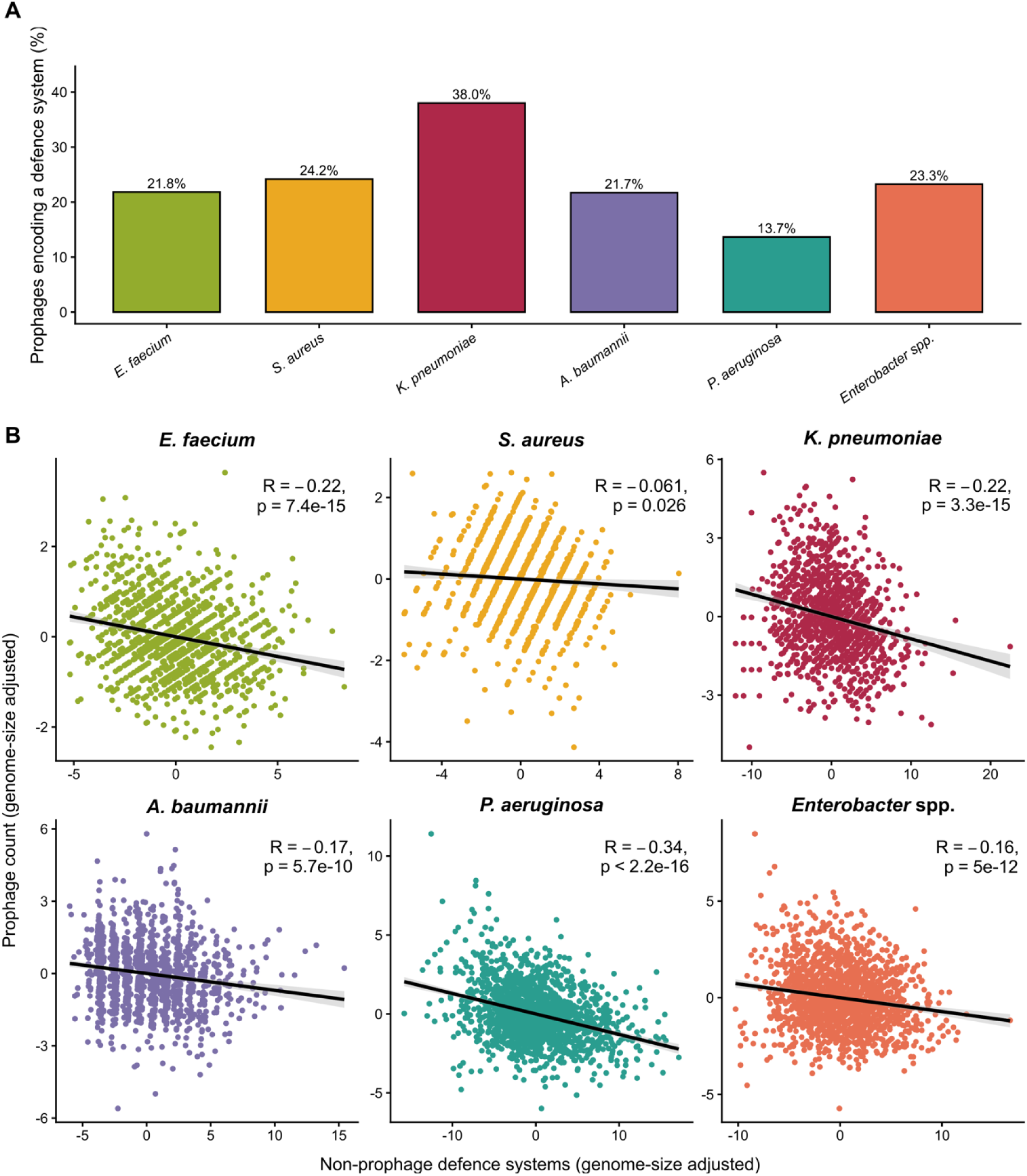
Defence Systems of ESKAPE Pathogens. **(A)** The percentage of prophages encoding at least one anti-phage defence system, for each ESKAPE pathogen. **(B)** Relationship between non-prophage defence-system count and prophage count for each ESKAPE pathogen, with both variables adjusted for bacterial genome length. Each axis shows the residual after regression on genome length, so the trend down corresponds to the partial correlation controlling for genome length. The black line shows the linear regression with its 95% confidence interval, and the Pearson coefficient and p-value are given for each pathogen.

### Non-prophage defence carriage is inversely associated with prophage carriage after accounting for genome length

Prophage count increases with bacterial genome length (above), and we found that non-prophage defence-system count also increased with genome length in all six pathogens (Pearson R = 0.14 to 0.71, all p < 0.01; Supplementary Figure 11). Genome length is therefore a shared correlate of both traits and confounds their direct comparison. Controlling for genome length, non-prophage defence-system count and prophage count were negatively correlated in every pathogen (partial R = −0.06 to −0.34, Figure 6B). This was significant in all six, although only marginally in *S. aureus* (R = −0.06, p = 0.03, compared with p < 0.001 in the other five). Genomes carrying more non-prophage defence systems thus tended to carry slightly fewer prophages than other genomes of similar size. The relationship was consistent across all six pathogens but weak in magnitude, and opposite in sign to the unadjusted trend.

To identify which systems contributed to this inverse relationship, we tested each defence subtype individually. For every subtype carried by between 5% and 95% of a pathogen’s genomes, we compared prophage counts between the genomes that did and did not carry that subtype, while accounting for genome length and for the remainder of each genome’s non-prophage defence repertoire. This approach isolated the association of an individual subtype with prophage carriage from the genome-wide trend described above. Of 203 pathogen and subtype combinations tested, 67 were significant after correction for multiple testing, and associations with reduced prophage carriage (n=46; Supplementary Figure 12) outnumbered associations with increased prophage carriage (n=21; Supplementary Figure 12). The subtypes linked to reduced prophage carriage included several well-characterised interference systems, among them the CRISPR-Cas subtypes IV-A and I-F, the abortive infection system AbiQ, CBASS, and Mokosh. Several of the subtypes linked to increased prophage carriage, including PD-T7-1, PARIS, and DS-24, are themselves frequently prophage-encoded, so their non-prophage copies may reflect a genomic background rich in mobile elements rather than a tendency to promote prophage carriage.

## DISCUSSION

PHORAGER is a scalable, modular pipeline for the identification, annotation, and quality assessment of prophages from bacterial genomes. The pipeline was designed to simplify prophage mining by only requiring a single fasta file or directory as input and by producing ready-to-analyse tsv summary tables, making large-scale prophage analysis more accessible to users without extensive bioinformatics expertise.

Application of PHORAGER to a large collection of publicly available ESKAPE pathogen genomes demonstrated that the pipeline can efficiently process diverse bacterial datasets, spanning both Gram-positive and Gram-negative organisms, and detect a broad range of prophages, including both double-stranded and single-stranded DNA forms. Our analysis of 30,824 genomes also represents the largest-scale prophage mining effort of the ESKAPE pathogens to date, including over 29,000 more genomes than previous studies. These studies by Botelho *et al*. (2023) and Kondo *et al*. (2021) analysed 1782 and 1623 genomes respectively, each relying on singular mining tools, phispy and PHASTER, with no downstream quality assessments performed (68,69). Beyond scale, these studies also differ in focus; Botelho *et al*. examined the complete ESKAPE mobilome, including plasmids and integrative conjugative elements, while Kondo *et al*. concentrated on the AMR and virulence factor genes encoded by the ESKAPE prophages (68,69). In contrast, our study introduces PHORAGER, emphasising methodological rigour in prophage recovery alongside offering biological insights into the ESKAPE prophage repertoire.

Our analysis showed that the majority of ESKAPE pathogens (93.6%) were lysogens harbouring one or more prophages, supporting the view that lysogeny is widespread among clinically relevant bacterial pathogens. We also observed a significant positive correlation between bacterial genome length and number of encoded prophages, corroborating previous studies suggesting that larger genomes provide greater opportunities for prophage acquisition and carriage (70).

The large-scale analysis also highlighted substantial levels of redundancy within publicly available databases. This was particularly evident for *S. aureus*, where dereplication at the strain level removed the majority of genomes, likely reflecting the repeated sequencing of successful epidemic lineages such as methicillin-resistant *S. aureus* (MRSA). In contrast, the greater genomic diversity observed in *Enterobacter* spp. was complemented by a larger and more diverse prophage repertoire, indicating that host diversity contributes to the range of prophage populations that can be recovered.

One of the distinguishing features of PHORAGER compared with existing prophage mining pipelines is the consolidation of overlapping predictions generated by multiple mining tools. Merging these overlapping sequences greatly reduces redundancy within the final prophage dataset and, in our analysis, increased the number of high-quality and complete prophage sequences recovered. The use of both VIBRANT and geNomad is still justified by the observation that, despite a considerable degree of overlap, each tool generated unique predictions, due to their differing approaches to viral sequence identification. The selection of VIBRANT and geNomad over alternative prophage mining tools was also intentional. Both combine sequence similarity-based methods with machine learning approaches, enabling the identification of both previously characterised and potentially novel prophages. Additionally, both tools are available on the command line, making them suitable for large-scale, high-throughput analyses. Web-based platforms such as PHASTEST offer excellent accessibility and ease of use, but are less practical for processing large genomic datasets (23). By integrating complementary prediction methods with a consolidation step, PHORAGER improves the recovery of high-quality prophages while minimising redundancy.

A second key feature of PHORAGER is the annotation-based filtering step implemented after CheckV quality assessment. This step was introduced after observing that some sequences classified by CheckV as medium-quality or above lacked characteristics typically associated with phages when their gene content was examined manually. Although CheckV is one of the most widely used tools for viral quality assessment, it assumes that the input sequences are confirmed as viral and estimates completeness and quality based on this premise. Consequently, false-positive predictions generated by prophage mining tools can be assigned high quality scores and be inadvertently retained in downstream analysis. Annotation-based filtering helps to address this limitation by incorporating functional evidence from Pharokka and Phold which are specifically designed for phage annotation. In the ESKAPE dataset, most sequences removed by this step had been assigned medium-quality by CheckV, however a small proportion had been classified as complete. Many of these ‘complete’ sequences were identified through high-confidence direct terminal repeats (DTRs), which in some cases can arise from short-read assembly artefacts or the presence of randomly repeated sequences in the bacterial genome, while others carried CheckV warnings indicating that no viral genes had been detected. These findings highlight the importance of integrating multiple forms of quality control to reduce false positives and improve the reliability of prophage datasets.

Five viral classes were identified in the final prophage dataset, spanning both double-stranded and single-stranded DNA phages. Nevertheless, the dataset was overwhelmingly dominated by double-stranded DNA phages assigned to the class *Caudoviricetes*. This is consistent with the current understanding that lysogenic double-stranded DNA tailed phages are the predominant form of prophages found within bacterial genomes, although this observation could also be influenced by biases in the reference databases used by many of the tools implemented by PHORAGER, or by the filtering thresholds selected. Analysis of the predicted viral genera indicated a high degree of host-specificity. However, where genera were shared between hosts, prophages from *K. pneumoniae* and *Enterobacter* spp. co-occurred most frequently, likely due to the close evolutionary relationship between these organisms.

Recent work by Perfilyev *et al*. (2026) identified thousands of bacterial assembly-associated phage sequences (BAPS), representing lytic phage genomes that can be recovered from bacterial sequencing data and potentially be falsely identified as prophages by mining tools (71). Certain viral taxa, including the newly proposed “*Bapsvirus*” genus within the *Chimalliviridae* family, appear to be strongly associated with this phenomenon. No members of this family or genus were detected in our final dataset, suggesting that PHORAGER may be relatively robust to this source of false positives. However, this does not exclude the possibility that lytic phages remain within the final prophage collection, and additional approaches for identifying and removing BAPS could be incorporated into future versions of the pipeline.

Screening of the final prophage dataset for putative ARGs and virulence factors demonstrated that prophages from all ESKAPE pathogens have the potential to contribute to bacterial pathogenicity. Putative ARG counts were generally low across the dataset, however *A. baumannii* prophages encoded the greatest number. This contrasts with the findings of Kondo *et al*. who reported that *K. pneumoniae* prophages encoded the largest repertoire of ARGs among the ESKAPE pathogens (69). Nonetheless, our observations for virulence factors were largely consistent with those of Kondo *et al*., with *S. aureus* prophages encoding by far the greatest number, while *A. baumannii* prophages had the fewest (69). These findings should however be interpreted with caution as *in silico* predictions of ARGs and virulence factors do not necessarily indicate functional activity (72). Experimental *in vitro* validation is therefore required before these genes can be confirmed to contribute to bacterial pathogenicity and AMR.

We also found that anti-phage defence systems were a common feature of the ESKAPE prophages, with 23.6% encoding at least one, identifying prophages as an important reservoir of bacterial immunity. Prophage-encoded defences are thought to protect the lysogen, and the resident prophage itself, against infection by competing phages through cognate self-immunity, a form of superinfection exclusion (8). Such exclusion could both stabilise existing lysogens and influence which further prophages a genome is able to acquire. Consistent with this, genomes encoding more non-prophage defence systems tended to carry slightly fewer prophages once genome length was taken into account, in keeping with anti-phage defences limiting phage acquisition. However, this association was weak and correlative, and several of the systems involved are themselves frequently prophage-encoded, so disentangling cause from genomic background will require experimental validation.

Despite its advantages, PHORAGER has several limitations. Due to the number of bioinformatics tools and reference databases it incorporates, a complete installation requires approximately 38.6 GB of storage. This requirement may be reduced if users have previously installed some of the required databases, however the overall storage requirement remains relatively large compared with simpler prophage mining workflows. A second limitation is that PHORAGER involves several filtering steps that require the user to select appropriate parameters for their dataset. Choosing suitable thresholds is vital but can be challenging, particularly for non-expert users, and no filtering strategy can completely eliminate the trade-off between sensitivity and specificity. Some false positives are likely to remain, and some true positives may be discarded. For example, PHORAGER’s default CheckV filtering criteria discards sequences classified as low-quality and ‘not determined’. Although this approach may exclude a small number of highly novel prophages, it was chosen to maximise the specificity of the final dataset. Users interested in discovering completely novel phages may therefore wish to investigate sequences assigned to the ‘not determined’ category.

Parameter selection becomes more nuanced during the annotation-based filtering step using Pharokka and Phold, where the objective is to discern viral from non-viral sequences based on gene content. Our analysis demonstrated that the default filtering thresholds remove a proportion of likely false positives. These thresholds were selected based on previous extensive manual inspection of annotated prophage genome plots, and we recommend that users also perform similar assessments when analysing new datasets to optimise filtering performance. However, we appreciate that not all users will be confident selecting custom thresholds, and therefore PHORAGER provides sensible default parameters that offer a practical starting point, though it may not be optimal for every application.

A broader challenge in large-scale prophage mining is the difficulty of benchmarking predictions because the true prophage content of bacterial genomes is rarely known with certainty. Even when extensive experimental validation is performed through induction assays, such as those reported by Dahlman *et al*., (2025), bioinformatically predicted prophages can remain difficult to validate (73). Not all prophages are inducible under the conditions tested, some might require highly specific induction triggers, and others may be cryptic prophages that have lost the ability to excise from the bacterial chromosome. Consequently, distinguishing true prophages from non-viral genomic regions remains a significant challenge in the field, and no current bioinformatics approach can establish a definitive ground truth. Although PHORAGER implements multiple prediction, quality control, and filtering steps to try and increase the reliability of the final prophage dataset, the recovered sequences should be treated as putative prophages pending experimental validation.

In conclusion, PHORAGER is an automated, scalable pipeline, developed to recover high-quality prophages from bacterial genomes. PHORAGER was designed to help standardise the process of prophage mining, providing suggestions on how best to quality check and filter both the bacterial inputs and putative prophages to ensure the final dataset contains limited false positives. We hope that users will be able to implement the pipeline into their workflows to easily analyse their own datasets and improve our collective understanding of prophage carriage rates and influence across different bacterial species.

## ACKNOWLEDGEMENTS

AUTHOR CONTRIBUTIONS

Xena Dyball: Conceptualisation, Methodology, Investigation, Visualisation, Writing-original draft. Alise J Ponsero: Data curation, Software, Formal analysis, Writing – review & editing. James A D Docherty: Conceptualisation, Methodology, Writing – review & editing. Andrea Telatin: Visualisation, Writing – review & editing. Emmanuelle H Crost: Supervision, Writing – review & editing. Nathalie Juge: Supervision, Writing – review & editing. Ryan Cook: Conceptualisation, Investigation, Visualisation, Supervision, Writing – review & editing. Evelien M Adriaenssens: Conceptualisation, Supervision, Writing – review & editing.

## DATA AVAILABILITY

PHORAGER is freely available as open-source software on Github (https://github.com/aponsero/PHORAGER) and workflow hub (PHORAGER v0.5.0). The ESKAPE prophage dataset is available on Zenodo (DOI:10.5281/zenodo.18328113).

## SUPPLEMENTARY DATA

**Supplementary figure 1:**
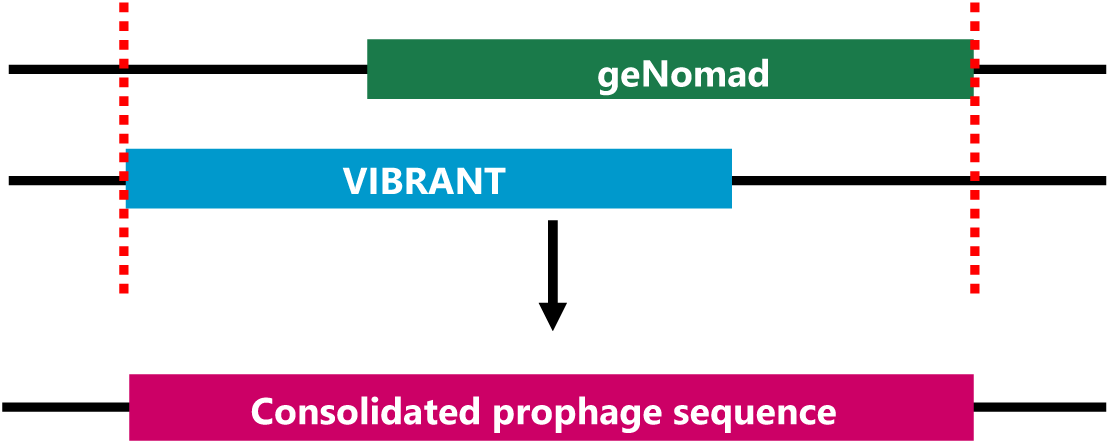
Coordinate consolidation rationale. An example of how the mining tools may identify different boundaries for the same prophage. In these instances, PHORAGER detects the overlapping regions and identifies the earliest start coordinate and latest end coordinate to produce a single consolidated prophage sequence, which is then used for the prophage processing module of the pipeline.

**Supplementary figure 2:**
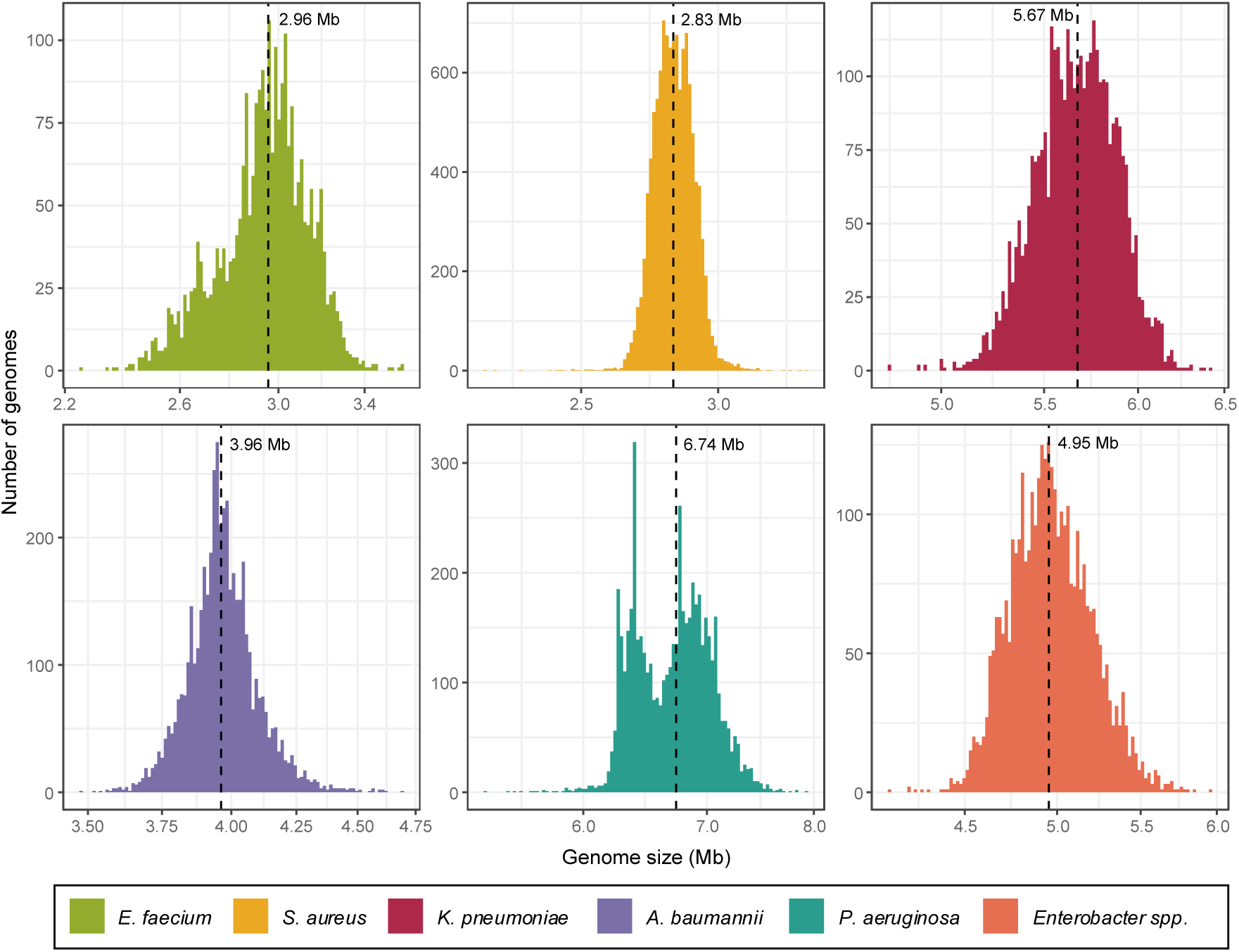
Histogram of ESKAPE pathogen genomes sizes. Distribution of bacterial genome lengths across the ESKAPE pathogens. The dashed black line indicates the median genome size for each pathogen.

**Supplementary figure 3:**
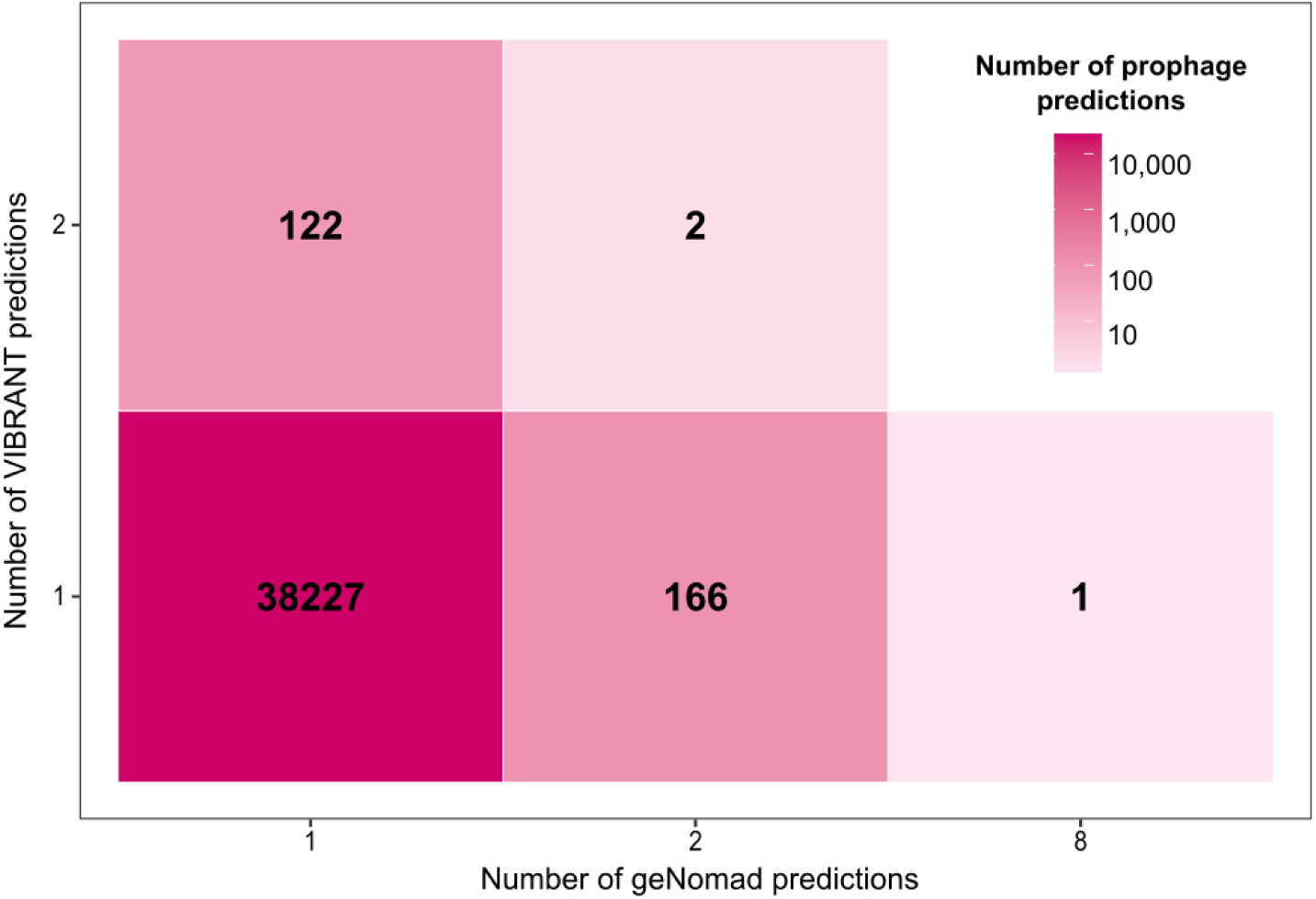
Tool prediction combinations underlying prophage consolidation. Heatmap of the different combinations of tool predictions merged in the consolidation step in PHORAGER prophage workflow.

**Supplementary figure 4:**
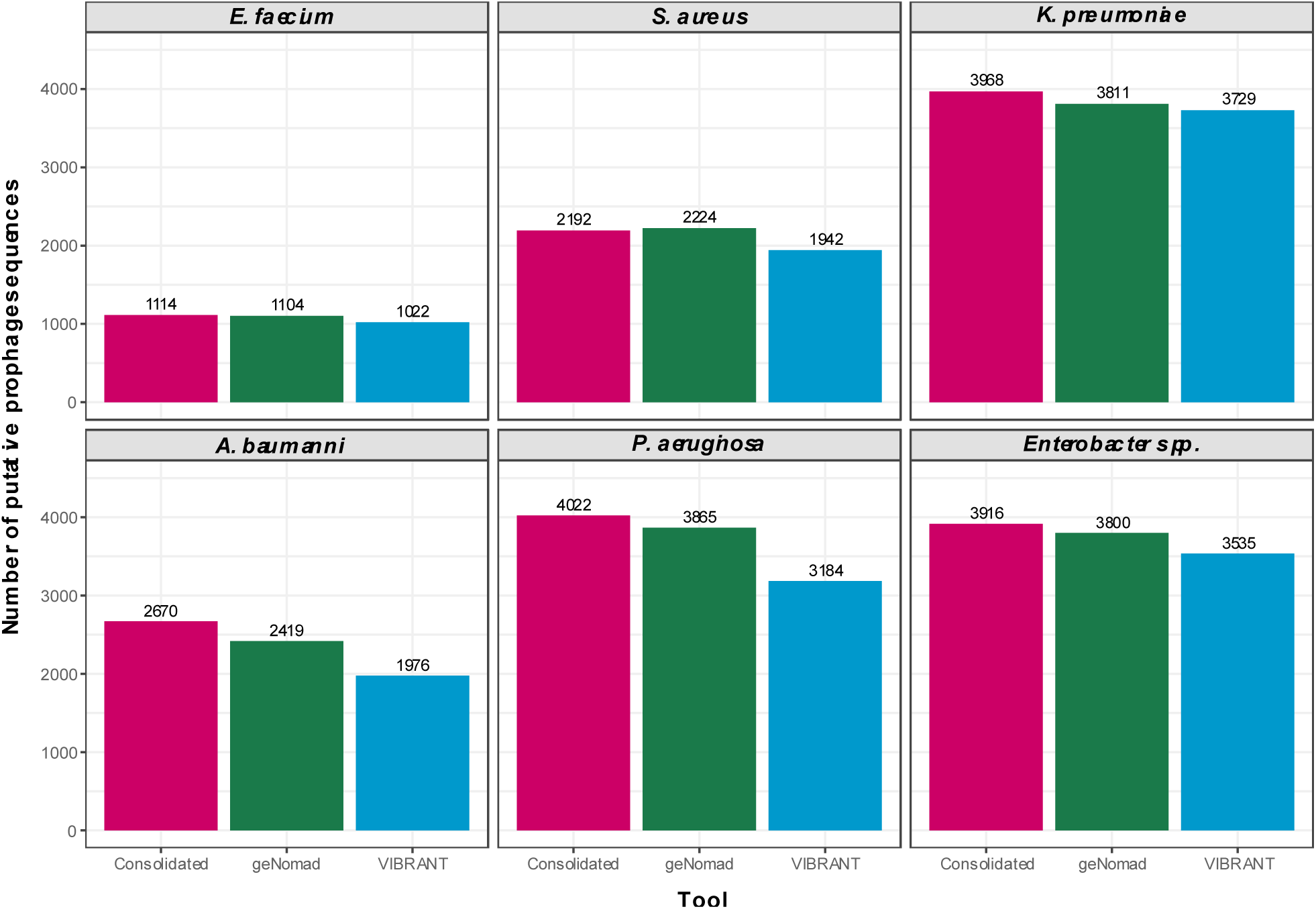
High-quality and complete prophage sequence counts for each tool. The number of putative prophage sequences assigned ‘high-quality’ or ‘complete’ by CheckV for the individual prophage mining tool predictions versus the consolidated sequences for each ESKAPE pathogen.

**Supplementary figure 5:**
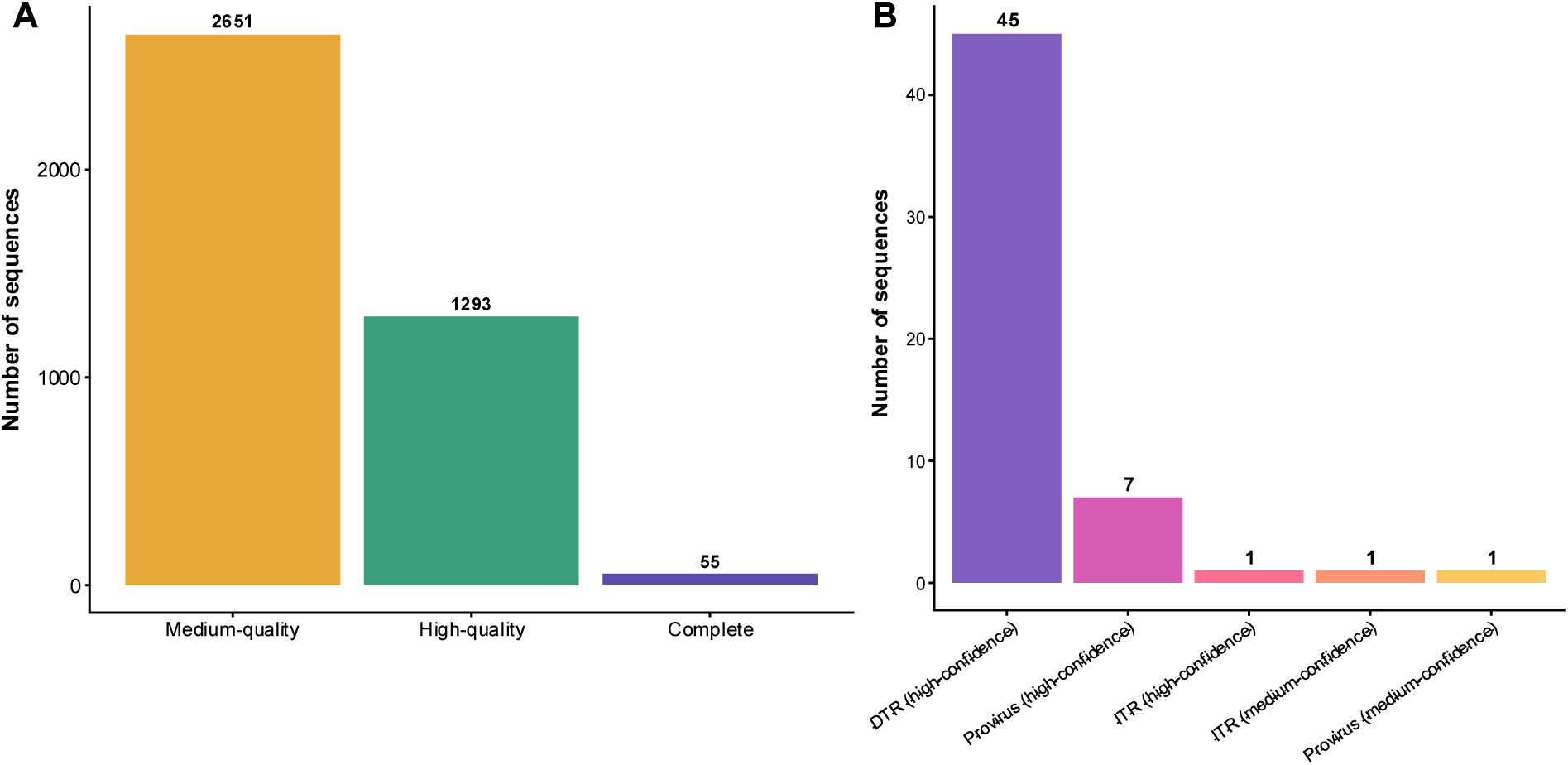
Quality scores and completeness methods for sequences removed by annotation-based sequences. **(A)** CheckV quality scores of the putative prophage sequences removed by annotation-based filtering during PHORAGER’s annotation workflow. **(B)** The completeness method used by CheckV for the ‘complete’ sequences removed by annotation-based filtering.

**Supplementary figure 6:**
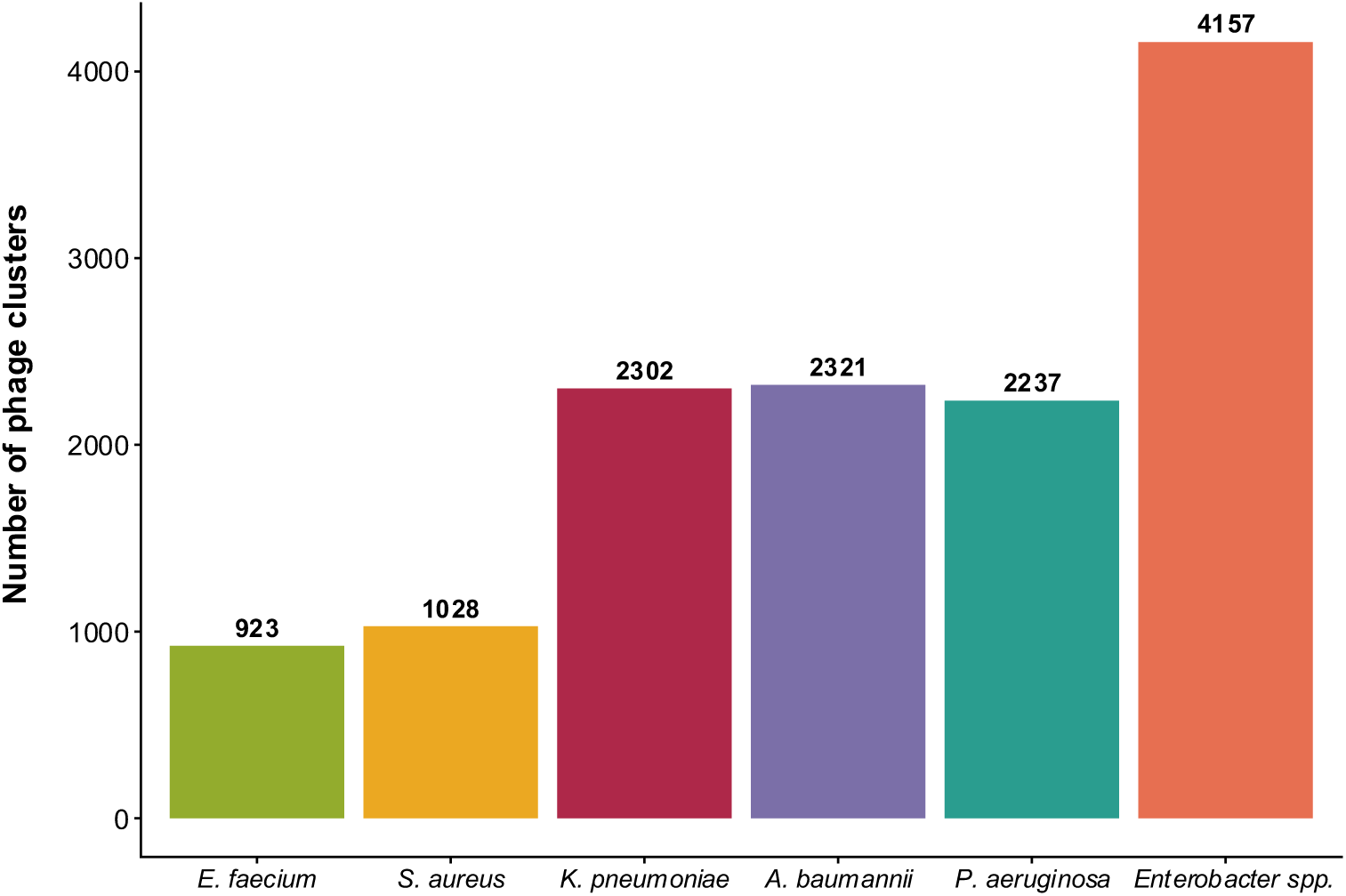
Number of phage clusters (vOTUs) identified for each ESKAPE pathogen. vOTUs were generated following dereplication at the approximate species level.

**Supplementary figure 7:**
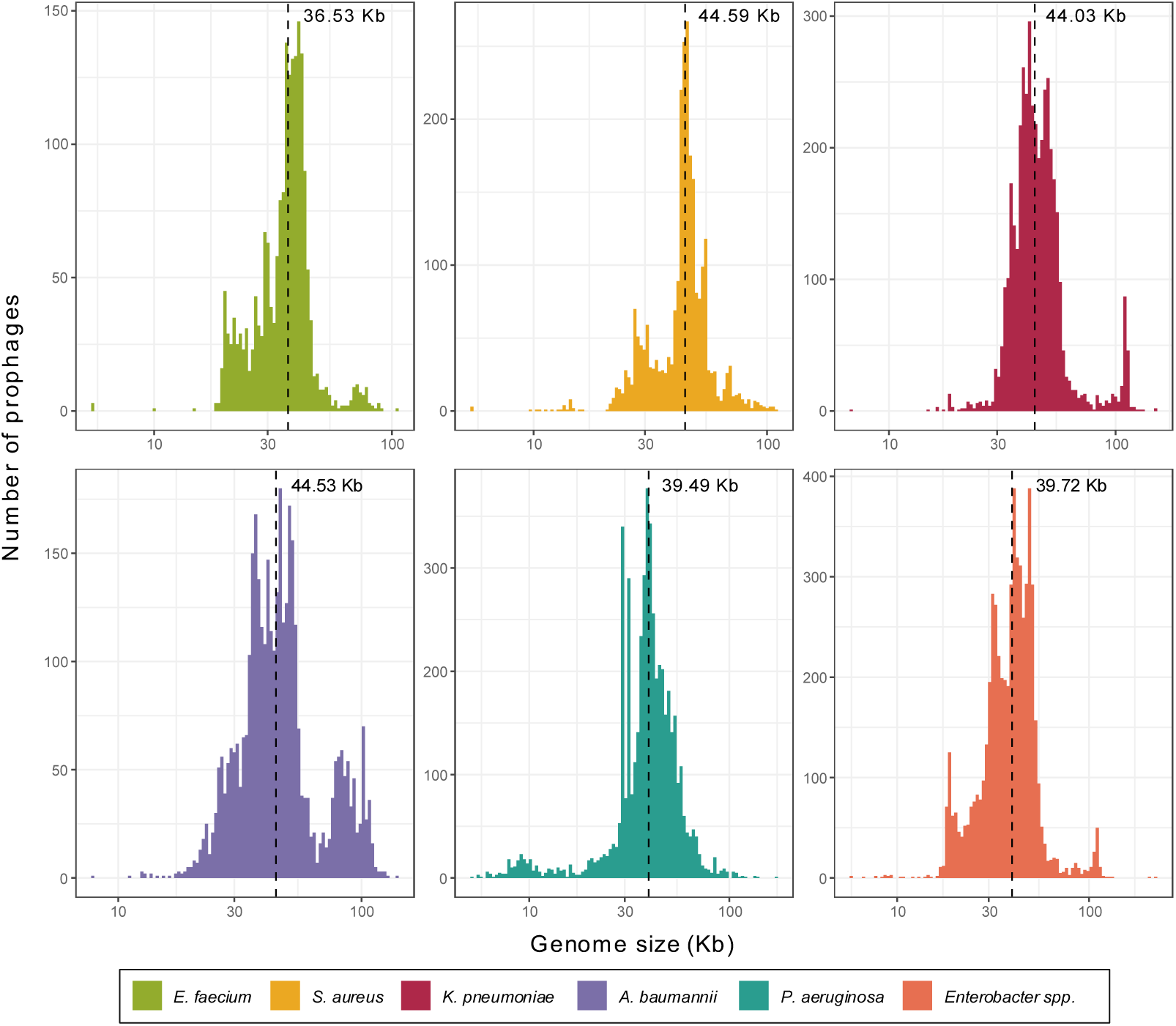
Prophage genome lengths. Distribution of prophage lengths across the ESKAPE pathogens. The dashed black line indicates the median prophage genome length.

**Supplementary figure 8:**
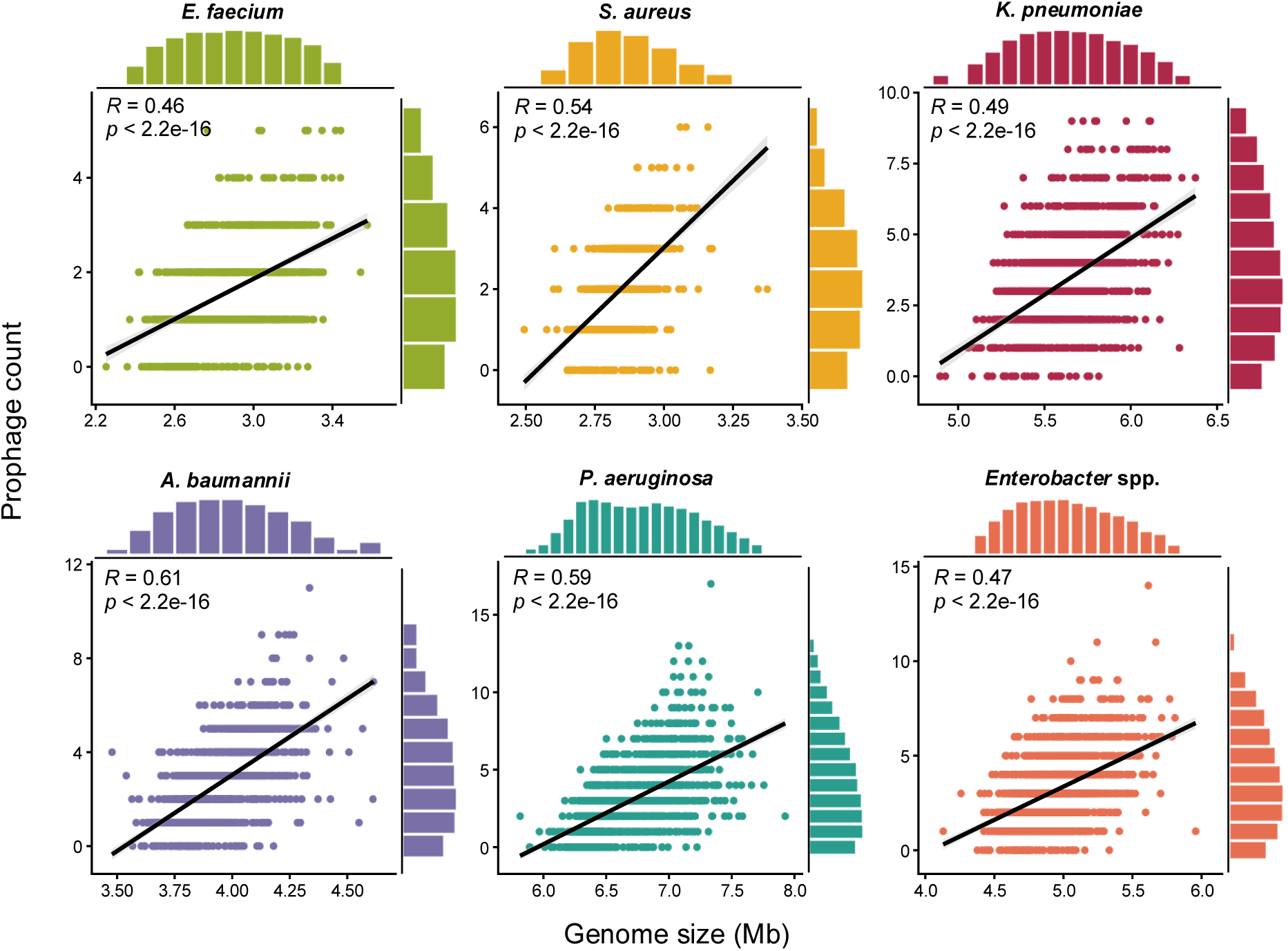
Genome length correlates with prophage count across ESKAPE pathogens. Correlation between bacterial genome length and prophage count for each ESKAPE pathogen.

**Supplementary figure 9:**
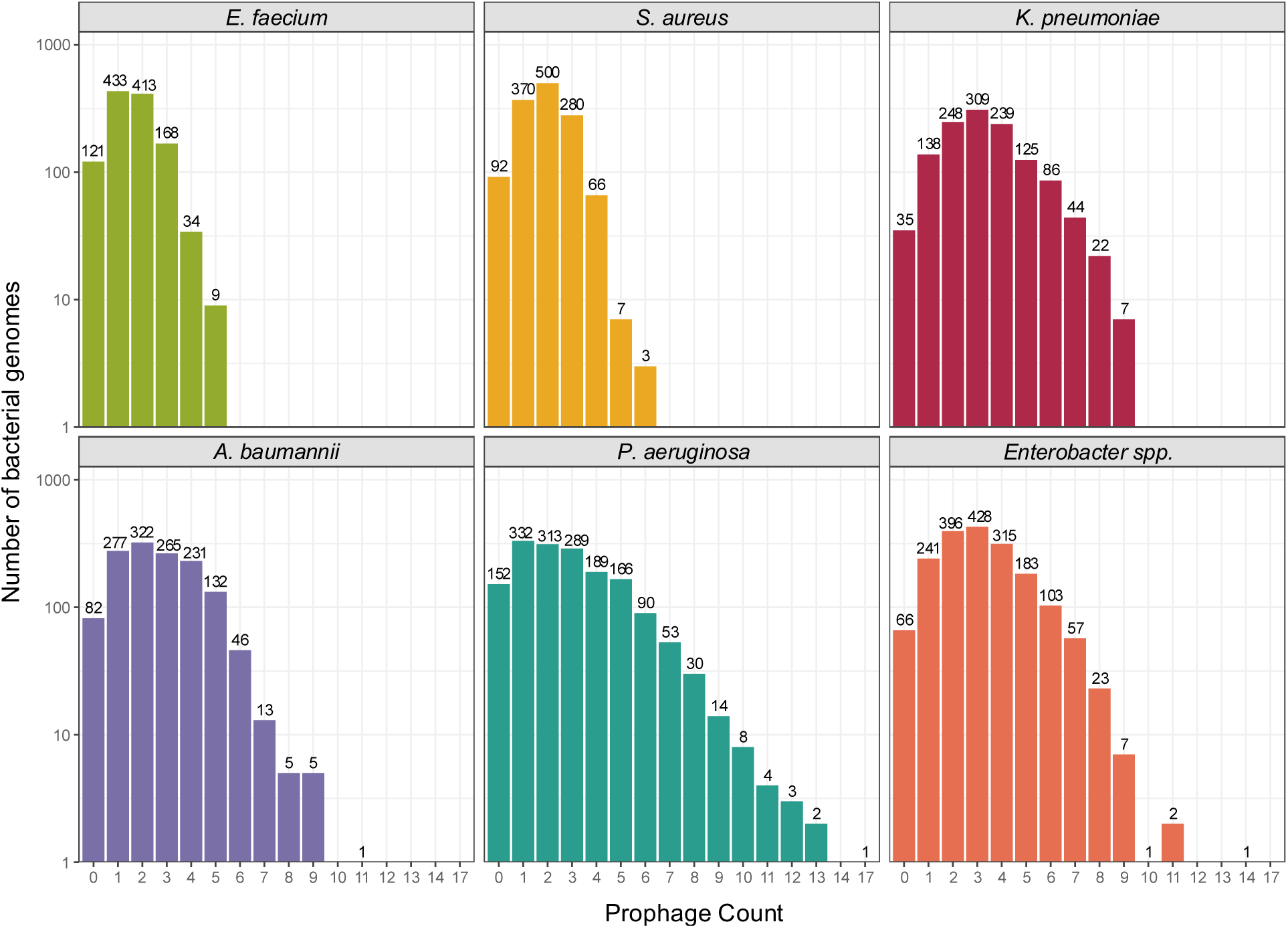
Prophage carriage among the ESKAPE pathogens. The number of bacterial genomes that have specific prophage counts across the ESKAPE pathogens.

**Supplementary figure 10:**
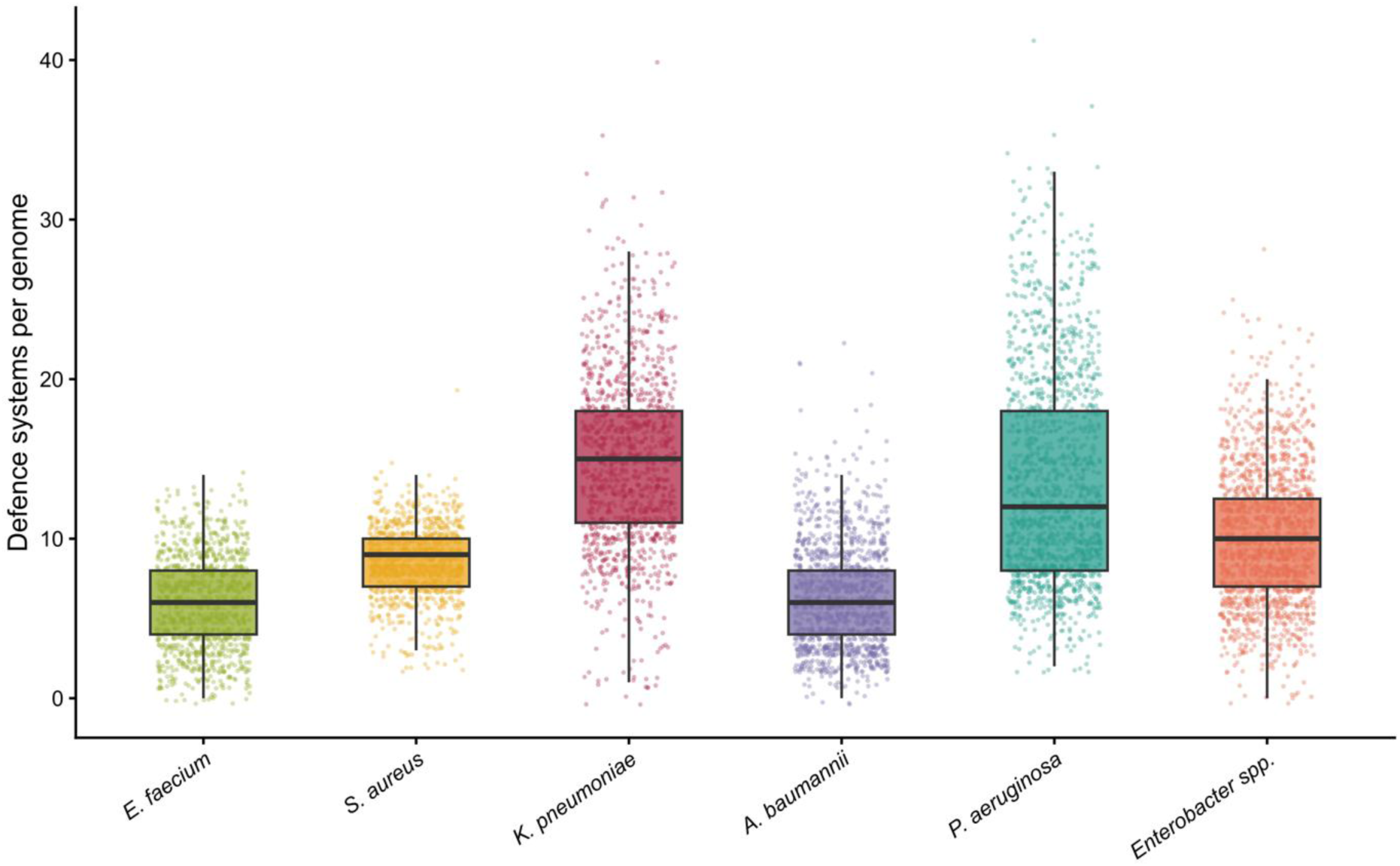
Defence systems per bacterial genome. The number of anti-phage defence systems encoded per genome across the ESKAPE pathogens. Boxes indicate the median and interquartile range, and points represent individual genomes.

**Supplementary figure 11:**
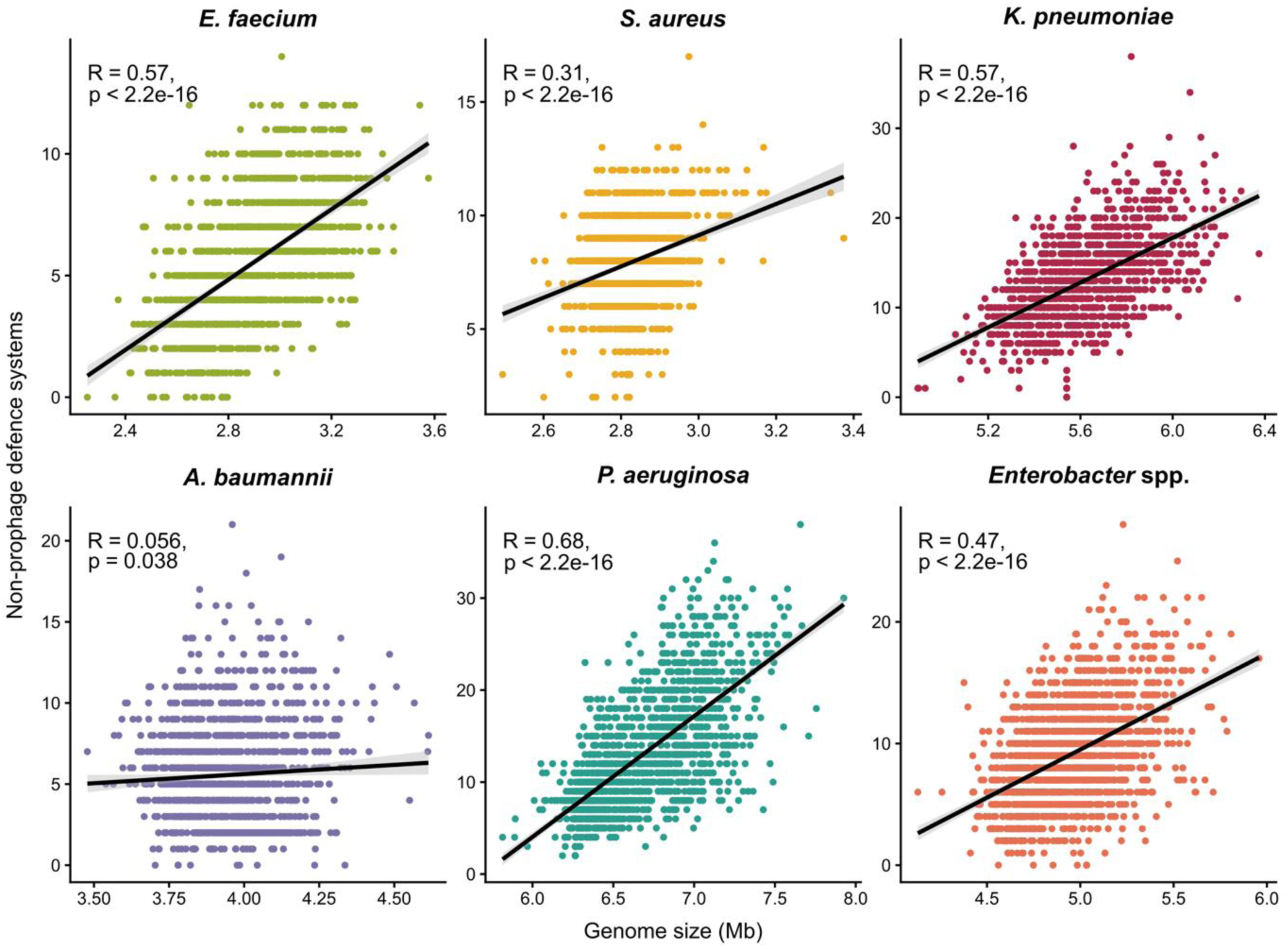
Genome length positively correlates with defence system carriage. Correlation between bacterial genome length and the number of non-prophage defence systems for each ESKAPE pathogen. The black line shows the linear regression with its 95% confidence interval, and the Pearson correlation coefficient and p-value are given for each pathogen.

**Supplementary figure 12:**
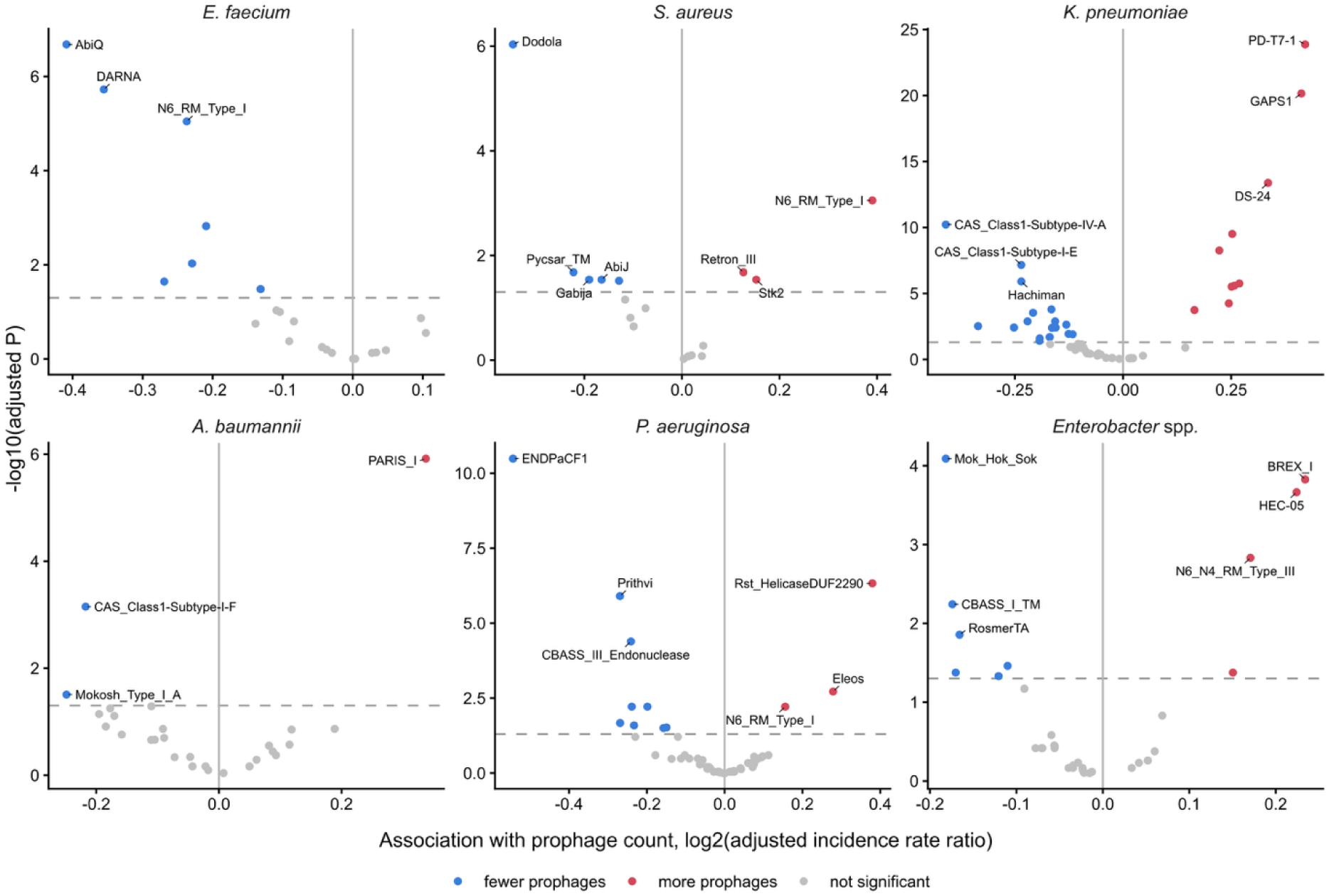
Association between carriage of individual non-prophage defence subtypes and prophage count, for each ESKAPE pathogen. Each point is a defence subtype; the x-axis shows the log2 adjusted incidence rate ratio (its effect on prophage count, adjusted for genome length and the rest of the non-prophage defence repertoire) and the y-axis shows the significance. Subtypes significantly associated with fewer or more prophages (Benjamini-Hochberg adjusted p < 0.05) are coloured blue and red respectively, with the three most significant in each direction labelled. The dashed line marks the significance threshold.

